# The unfolded protein response switches from pro-survival to pro-apoptotic to cap the extent of endoplasmic reticulum expansion

**DOI:** 10.1101/134726

**Authors:** Milena Vitale, Andrea Orsi, Anush Bakunts, Federica Lari, Laura Tadè, Angela Cattaneo, Roberto Sitia, Andrea Raimondi, Angela Bachi, Eelco van Anken

## Abstract

Insufficient folding capacity of the endoplasmic reticulum (ER) activates the unfolded protein response (UPR) to restore homeostasis. Yet, how the UPR achieves and evaluates ER homeostatic readjustment is poorly understood. In a HeLa cell model we show that, upon a severe proteostatic insult that eclipses the ER chaperone BiP, the UPR transitions from acute full-gear activation to chronic submaximal activation, when BiP is in excess again. As such, the UPR-driven ER expansion is pro-survival, primarily via the UPR transducer ATF6α. Simultaneous abrogation of ER-associated degradation (ERAD) leads to chronic full-gear UPR activation and further ER expansion, but the UPR transducers IRE1α and PERK turn pro-apoptotic. Pro-survival *XBP1* mRNA processing by IRE1α is then trumped since unspliced *XBP1* mRNA is depleted and IRE1α’s endonuclease activity is unleashed against other RNAs. Thus, the IRE1α/*XBP1* relay serves as a capacitor of ER expansion that heralds cell death if the expansion is deemed excessive.

## INTRODUCTION

Key to survival is that organisms adapt their behavior—or on the cellular level their molecular machineries—according to need. An example is how the endoplasmic reticulum (ER) accommodates fluctuations in the load of client proteins that fold and assemble there before being dispatched to travel further along the secretory pathway (***Walter and Ron, 2011***). An impressive array of ER resident chaperones and other folding factors assist the maturation of client proteins (***van Anken and Braakman, 2005***). BiP is an ER resident HSP70 family chaperone, whose function is regulated by at least five different HSP40 co-chaperones, two nucleotide exchange factors, and through reversible AMPylation (***Preissler et al., 2015***). The ER also hosts an HSP90 family member, GRP94, for which CNPY family proteins may be co-chaperones (***Anelli and van Anken, 2013***), and several PPIases, such as cyclophilin B and a few FKBP family members. Special to the ER are the chaperones calnexin (CNX) and calreticulin (CRT) that bind N-glycans on client proteins, and numerous protein disulfide isomerases (PDI) that assist disulfide bond formation (***van Anken and Braakman, 2005***).

The chaperones and folding factors retain folding and assembly intermediates in the ER so as to minimize the secretion or deployment throughout the endomembrane system of immature client proteins (***Ellgaard and Helenius. 2003***). Such client retention in the ER implies that when the protein folding machinery is inadequate in letting clients mature, they accumulate, which causes ER stress and activates an adaptive response: the UPR (***Walter and Ron, 2011***). The circuitry of the UPR has been mapped to impressive detail: the three well-studied branches of the UPR are initiated by the UPR transducers IRE1α, PERK and ATF6α, each having an ER stress sensing domain in the ER lumen and a UPR signaling effector domain in the cytosol. Activated IRE1α and PERK are kinases that trans-autophosphorylate. This step triggers IRE1α to assume endonuclease activity such that it removes an intron from *XBP1* mRNA. Upon its religation the spliced mRNA encodes the XBP1 transcription factor (***Yoshida et al., 2001; Calfon et al., 2002***). The IRE1α endonuclease activity may also stray to other RNAs in a process named regulated IRE1-dependent decay (RIDD; ***Hollien and Weissman, 2006; Hollien et al., 2009***). Activated PERK transiently attenuates protein synthesis through phosphorylation of the translation initiation factor eIF2α (***Harding et al., 1999***). At the same time eIF2α phosphorylation favors expression of a few transcripts, in particular ATF4 and ATF5, which are transcription factors that activate further downstream effectors, such as CHOP (***Walter and Ron, 2011***). The third UPR branch is activated by ATF6α that undergoes regulated intramembrane proteolysis in the Golgi, which liberates a transcriptionally active N-terminal portion of 50 kDa that acts as a transcription factor (***Ye et al., 2000***). The UPR-driven mobilization of these transcription factors ensures that through their concerted efforts a comprehensive genetic program is initiated that drives expression of all components that are necessary to expand the ER, including chaperones and enzymes for membrane synthesis (***Walter and Ron, 2011***). In fact, overexpression of for instance XBP1 alone leads to ER expansion even in absence of any perturbation of the ER client protein folding and assembly process (***Sriburi et al., 2000***). Moreover, UPR activation reinforces the ERAD machinery that dispatches ER clients that fail to fold or assemble across the ER membrane to the cytosol for proteasomal destruction (***Walter and Ron, 2011***).

Altogether, the UPR aims for homeostatic readjustment of the ER folding machinery through expansion of the organelle when needed, as well as for making life/death decisions depending on the severity of ER stress (***Walter and Ron, 2011***). Indeed, the UPR displays pro-apoptotic activity, which contributes to disease when dysregulated. For instance, UPR-driven apoptosis in β-cells of the pancreas seems to be a major cause of Type II diabetes (***Scheuner and Kaufman, 2008***) and various other pathologies (***Wang and Kaufman, 2012***). However, the question why the UPR would invoke pro-apoptotic pathways has been poorly addressed. Certainly, unbridled ER expansion occurs in so-called ER storage diseases (***Rutishauser and Spiess, 2002; Anelli and Sitia, 2010***) when ER clients accumulate but apparently no UPR-driven cell death is invoked. In epithelial cells that line the thick ascending limb of Henle’s loop in the nephron, ER accumulation of mutant uromodulin causes a UPR and such unrestrained expansion of the organelle (***Nasr et al., 2008***) that it leads to loss of cellular and tubule architecture (***Kemter et al., 2017***), and, ultimately, loss of kidney function (***Bernascone et al., 2010***). Along these lines, one could speculate that unrestrained ER expansion may cause gain-of-function disease irrespective of the ER client that drives the expansion. Pro-apoptotic output of the UPR could serve to afford cells self-censorship when they expand their ER to a size that is out of order.

Thus far, most studies on the UPR circuitry have focused on the signaling pathways themselves, while little is known about how the UPR evaluates the severity of ER stress and what defines the success of homeostatic readjustment of the ER. We here show that the widely used strategy of employing ER stress-eliciting drugs obscures how ER homeostatic readjustment may be achieved, and we present a HeLa cell model that instead allows to evaluate just that. By inducible overexpression of orphan immunoglobulin M (IgM) heavy chain (μ_s_), we provoke a full-blown UPR, which is essential for the cells to cope with the proteostatic insult. As μ_s_ accumulates in the ER, it temporarily reaches levels that are at least on a par with BiP, which provokes the UPR that lets BiP increase to levels that exceed μ_s_ levels again, while the ER expands in the process. The activation of the UPR is maximal only when there is a relative shortage of BiP, while it subsides to chronic submaximal output levels when ER homeostatic readjustment is achieved. Co-expression of Ig light chain (λ) instead leads to productive IgM secretion, such that BiP is not sequestered by μ_s_, the UPR is not activated and the ER does not expand. Conversely, abrogation of ERAD leads to further ER expansion since μ_s_ accumulates even more as it cannot be disposed of. Under these conditions IRE1α and PERK are chronically activated to maximal levels, which is when they relinquish their pro-survival role. While IRE1α still sustains maximal levels of *XBP1* mRNA splicing it also commits to RIDD. Apparently, IRE1α’s endonuclease activity is then unleashed against other RNAs, simply because levels of unspliced *XBP1* mRNA become limiting. As such, the IRE1α/*XBP1* relay serves as a capacitor that affords capping of ER expansion if it is judged to be excessive.

## RESULTS

### Cytotoxicity of UPR eliciting drugs

Drugs that typically are used to study the unfolded protein response (UPR) include tunicamycin (Tm), which prevents addition of N-glycans to nascent ER clients, dithiothreitol (DTT), which impedes disulfide bond formation, and thapsigargin (Tg), which depletes Ca^2+^ from the ER lumen. Their immediate effect is a general collapse of productive protein folding in the ER, and these drugs activate as such the UPR (***Walter and Ron, 2011***). In the longer run, however, these drugs likely have pleiotropic effects. Evidently, non-productive folding in the ER causes that ER clients are retained in the ER (***Ellgaard and Helenius, 2003***), and thus no longer reach their destination, be it anywhere throughout the endomembrane system of the cell or extracellularly. As a result—depending on their half-life (*t*½)—these proteins will be depleted at the site where they have to exert their functions, which ultimately may cause a plethora of perturbations to the homeostasis of the cell.

In order to unmask pleiotropic effects of UPR eliciting drugs, taking Tm as an example, we serially diluted it starting from a concentration (5 μg/ml) that is widely used to study the effects of ER stress and UPR activation, and monitored cell survival after 7 days by a cell growth / colony formation assay. At this high concentration a full-blown UPR was triggered, which is evident from the maximal extent of *XBP1* mRNA splicing in HeLa cells, i.e. the appearance of a band of higher mobility, corresponding to the RT-PCR product of the *XBP1^S^* transcript from which the intron has been removed (***Figure 1A***). Yet, there was no cell survival after 7 days (***Figure 1B***); not even when the cells were treated with Tm only the first day for 4 hrs. At a 100-fold dilution of the “standard” concentration, Tm marginally induced splicing of *XBP1* mRNA (***Figure 1A***), but still provoked a substantial cell growth defect (***Figure 1B,C***). Even at a further two-fold dilution of Tm to 25 ng/ml that failed to engender *XBP1* mRNA splicing altogether (***Figure 1A***) there was some loss of cell growth (***Figure 1B,C***). These findings thus reveal that an overall impairment of ER client protein folding has cytotoxic consequences aside from the activation of the UPR *per se*. Indeed, Tm readily causes morphological aberrations of the ER also at these low concentrations (***Rutkowski et al., 2006***).

**Figure 1.**
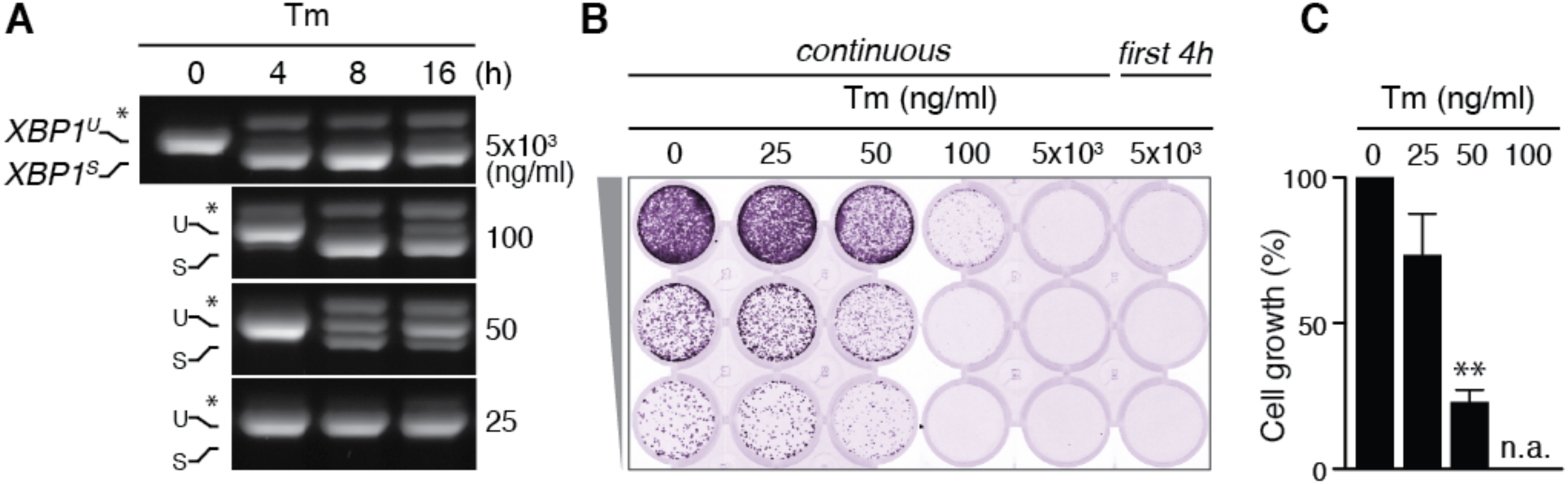
Tunicamycin has pleiotropic cytotoxic effects. (**A**) HeLa S3 cells were treated with Tm at various concentrations and for various durations, as indicated, before mRNA was isolated from cells for RT-PCR analysis with oligos specific for XBP1. PCR fragments corresponding to spliced (*XBP*^*S*^) and unspliced (*XBP*^*U*^) XBP1 were separated on gel. A hybrid product that is formed during the PCR reaction (***Shang and Lehrman, 2004***) is denoted by an asterisk. The ratio of (*XBP*^*S*^)/(*XBP*^*S*^ + *XBP*^*U*^) is indicative for activation of IRE1α. (B) Cells as in (**A**) were seeded upon 1:5 serial dilution into 24-well plates and treated continuously or only the first 4 hrs upon seeding with various concentrations of Tm, as indicated. After 7 days of growth, cells were fixed and stained with crystal violet. (**C**) Quantitation of crystal violet staining shown in (**B**) as a measure for cell growth, except for conditions that fully abrogated growth; n.a. = not assessed. Staining of untreated cells was set at 100%, and values are shown in bar graphs. Mean and s.e.m. are shown; n=3. Statistical significance in a one sample *t*-test of differences in crystal violet staining compared to untreated in (**B**) and replicate experiments was determined as a proxy for growth (**p ≤ 0.01).

### Proteostatic induction of the UPR as a model to assess ER homeostatic readjustment

We reasoned that pleiotropic effects of UPR eliciting drugs likely obscure important aspects of how cells cope with ER stress. If cell growth is compromised or, worse, if cell death is provoked already through these pleiotropic effects, how can we appreciate the contribution of the UPR to life/death decisions? Moreover, will UPR-driven ER expansion reach its full extent, if other cellular functions are compromised? We therefore sought to circumvent the shortcomings of using ER stress-eliciting drugs, and we hypothesized that over-expression of an ER client protein would be a better tool to appreciate these two aspects of UPR-driven adaptations to ER stress.

The impressive ER expansion during B cell differentiation (***Wiest et al., 1990; van Anken et al., 2003***) inspired us to choose the bulk product of plasma cells, secretory antibody, as an ideal candidate to engender a proteostatically driven UPR. More specifically, we decided to employ the secretory heavy chain (HC) of immunoglobulin M (μ_s_), since μ_s_ is expressed in bulk exclusively once B lymphocytes are activated and develop an expanded ER to become plasma cells (***Sitia et al., 1987***), and since plasma cell differentiation requires a functional UPR (***Reimold et al., 2001***). Yet, the precise role of the UPR in ER expansion and life/death decision in plasma cell development is difficult to assess. On the one hand, B cells anticipate that they will secrete IgM in bulk once they have differentiated into plasma cells (***van Anken et al., 2003***), such that ER expansion is also driven by developmental cues (***Iwakoshi et al., 2003***). Accordingly, ER expansion occurs in differentiating B cells even when μ_s_ is ablated (***Hu et al., 2009***). On the other hand, plasma cell life span is inherently limited out of immunological necessity, such that the UPR may contribute to life/death decisions in plasma cells, but unlikely exclusively so.

To assess the effect solely of μ_s_ expression on cellular homeostasis we thus decided to create a HeLa cell model that allowed inducible expression of murine μ_s_ for which we exploited the GeneSwitch system (Invitrogen). By subsequent lentiviral delivery and cloning of integrant cells we introduced the hybrid nuclear receptor that is activated by the synthetic steroid, mifepristone (Mif), yielding HeLa-MifON cells. The μ_s_ transgene under control of the promoter element responsive to that hybrid nuclear receptor was then integrated into the HeLa-MifON cells, yielding HeLa-μ_s_ cells.

Expression of μ_s_ in HeLa-μ_s_ cells in the absence of Mif was undetectable by immunoblotting (***Figure 2A***) or immunofluorescence (***Figure 2B***). The GeneSwitch system is amplified through an autoregulatory feedback loop whereby the Mif-bound hybrid nuclear receptor upregulates expression of its own gene. This feature likely causes that titration of μ_s_ expression levels by varying Mif concentration was not easily reproducible, and that the Mif-driven inducibility was rather an off/on effect (data not shown). A proxy for tunable expression of μ_s_ instead was offered by comparing different HeLa-μ_s_ clones, which—due to differences in genomic locus or copy number of integrations—each expressed μ_s_ at a different level upon induction with Mif as shown for three individual clones (***Figure 2A***). As illustrated by immunofluorescence of μ_s_ (***Figure 2B***), expression was remarkably uniform between cells since they were clonal. Moreover, μ_s_ is retained in the ER lumen, as shown by its strict co-localization with the ER marker CRT (***Figure 2C***).

**Figure 2.**
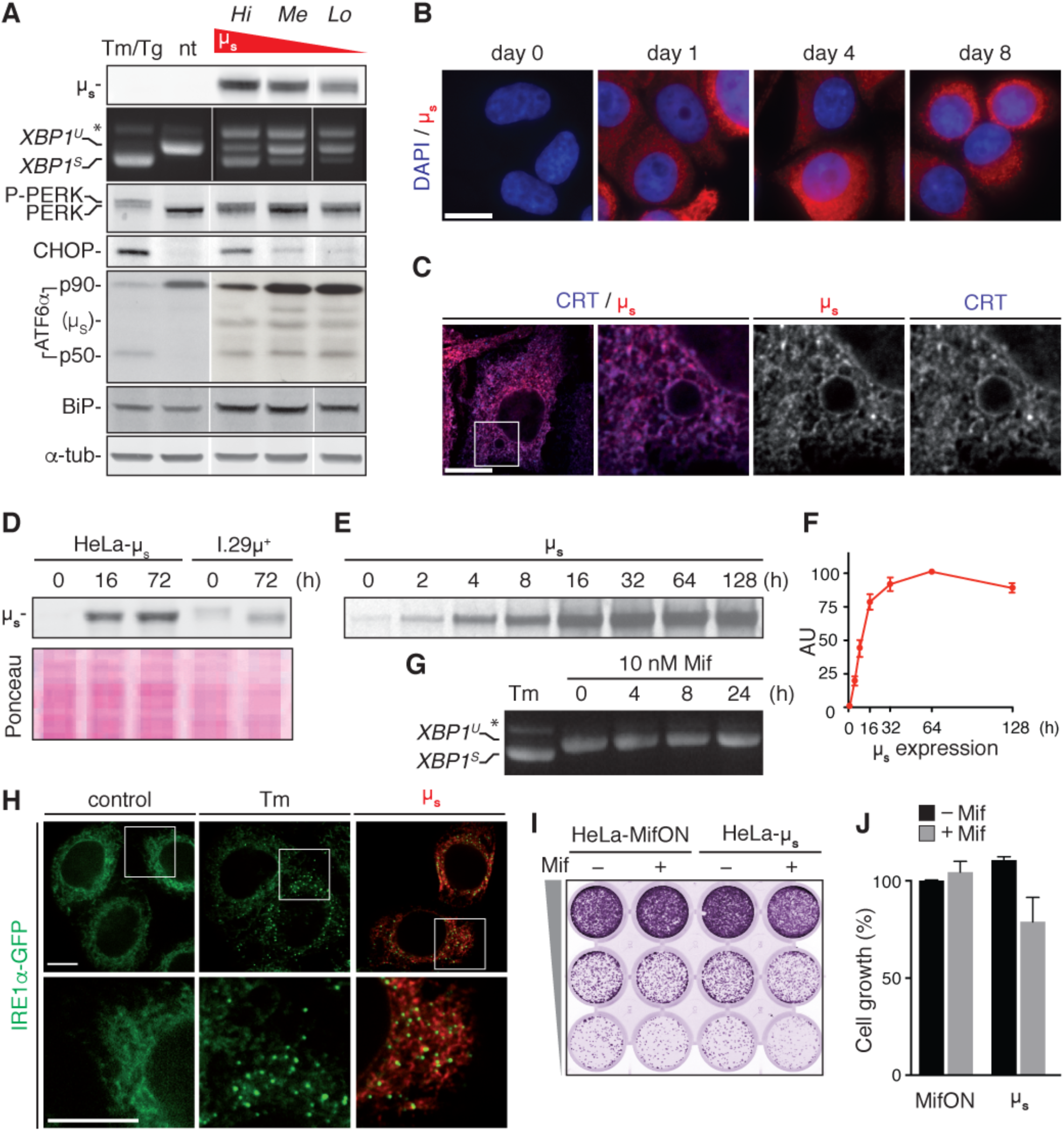
A model for proteostatically induced ER stress. (**A**-**F**,**H**-**J**) Expression of μ_s_ in HeLa-μ_s_ cells was induced with 0.5 nM Mif. (**A**,**G**,**H**) The UPR was pharmacologically induced as a reference for 4 hrs with 5 μg/ml Tm or for 1.5 hrs with 300 nM Tg—for analysis of ATF6α in (**A**) only, since Tm leads to its deglycosylation. (**A**,**D**) Immunoblotting revealed levels of μ_s_, CHOP, BiP and α-tubulin. A shift to a lower mobility phosphorylated form, P-PERK, and the appearance of CHOP revealed activation of PERK. The release of the p50 cleavage product from the p90 precursor revealed activation of ATF6α. Cross-reaction of the secondary antibody against anti-ATF6α with μ_s_ is denoted. (**A**,**G**) Splicing of XBP1 was assessed as in ***Figure 1A***. (**A**) Three different clones of HeLa-μ_s_ cells with decreasing μ_s_ expression levels: high (Hi), medium (Me), or low (Lo) were induced for 16 hrs. Non treated (nt) or Tm treated Hi HeLa-μ_s_ served as references. (**B**,**C**,**H**) Immunofluorescence revealed μ_s_ (red, **B**,**C**,**H**) and CRT (blue), which marks the ER (**C**) at the indicated times before or after induction (**B**) or after 8 hr induction (**C**,**H**). Nuclei were stained with DAPI (blue) (**B**). (**C**,**H**) The area that is boxed is shown by 3.5-fold magnification; scale bars represent 10 μm. (**D**) Samples were derived from equal numbers (7x10^4^) of HeLa-μ_s_ or I.29μ^+^ lymphomas induced with Mif, respectively, 20 μg/ml LPS for the indicated times. Ponceau staining of the blot serves as a loading control. (**E**) HeLa-μ_s_ cells were pulse labeled with 35S labeled methionine and cysteine for 10 min at the indicated times after induction. Immunoprecipitated μ_s_ was resolved by gel electrophoresis (**F**). Levels of radio-labeled μ_s_ in (**E**) were quantified by phosphor imaging. Maximal signal for μ_s_ (at 64 hrs) was set at a 100. (**G**) HeLa-MifON cells were treated with a high dose of Mif (10 nM) for the indicated times. (**H**) In HeLa-μ_s_ cells in which IRE1α was replaced with doxycycline (Dox) inducible IRE1α-GFP (green), its expression was tuned with 10 nM Dox to levels that allowed satisfactory detection of IRE1α-GFP by fluorescence microscopy. (**I**) Cell growth assay as ***Figure 1B*** of HeLa-MifON and HeLa-μ_s_ induced with Mif or not. (**J**) Quantitation of (**I**), performed as in ***Figure 1C***; Mean and s.e.m. are shown, n=2. There is no statistical significance in a one sample *t*-test of differences in growth between conditions.

The autoregulatory feedback loop of the GeneSwitch system has the advantage that it allows highly abundant expression of the transgene. Indeed, in the most highly expressing HeLa-μ_s_ clone μ_s_ reached levels that are at least on a par with those in model plasma cells: I.29μ^+^ lymphomas that were stimulated with lipopolysaccharide (LPS) for 3 days (***Figure 2D***). Synthesis of μ_s_ reached a plateau after 1-2 days of consistently high levels as judged by radiolabeling (***Figure 2E***,***F***). Thus, the proteostatic UPR stimulus is permanent, which implies that the cells must undergo homeostatic readjustment to meet the increased demand on the ER folding machinery.

By a series of standard assays, we then demonstrated that μ_s_ triggered a full-blown UPR. Upon 16 hrs of induction of μ_s_ expression, splicing of *XBP1* mRNA, phosphorylation of PERK (appearance of a band of slightly lower mobility), expression of PERK’s downstream effector CHOP, cleavage of the precursor ATF6α-p90 and the concomitant release of its cytosolic portion p50, were all at levels comparable to those triggered by a conventional Tm or Tg elicited UPR in these cells (***Figure 2A***). Treatment of the parental HeLa-MifON cells with a high dose of Mif (10 nM) did not provoke *XBP1* mRNA splicing (***Figure 2G***). Thus, the UPR induction in Mif treated HeLa-μ_s_ was the result of μ_s_ expression. Moreover, UPR activation upon μ_s_ induction was dose dependent, since the extent of *XBP1* mRNA splicing or CHOP expression was commensurate with the levels of μ_s_ that were expressed by the different HeLa-μ_s_ clones (***Figure 2A***). Another hallmark of a full-blown UPR activation is the clustering of IRE1α (***Li et al., 2010***). We deleted IRE1α by CRISPR/Cas9 and reconstituted it with GFP-tagged murine IRE1α (IRE1α-GFP) under control of a tight Tet responsive element (Clontech) by lentiviral transduction. The GFP-tag was introduced such that it did not interfere with function (***Figure 2—figure supplement 1***) as has been shown before for human IRE1α (***Li et al., 2010***) or yeast Ire1p (***Aragón et al., 2009***). With doxycycline (Dox) we tuned IRE1α-GFP expression to near-endogenous levels that allowed visualization by fluorescent microscopy and found that IRE1α-GFP not only clustered upon Tm treatment, but also upon μ_s_ expression (***Figure 2H***). Altogether, we concluded that μ_s_ expression drove activation of all branches of the UPR we tested in a similar manner and to a similar extent as triggered by ER-stress eliciting drugs.

Strikingly, in spite of the expression of μ_s_ in bulk, cell growth of the HeLa-μ_s_ cells almost paralleled that of the Mif treated parental MifON cells (***Figure 2I***,***J***). Thus, the full-blown UPR that was elicited by μ_s_ expression was well tolerated by the cells, unlike prolonged treatment with Tm at concentrations that triggered a full-blown UPR (***Figure 1***). In all, we concluded that the HeLa-μ_s_ cells offer a suitable model to assess ER homeostatic readjustment and the role of the UPR in the process since the cells apparently cope well with the proteostatic insult that is unabated once μ_s_ expression is induced.

### UPR activation correlates with accumulation of ER load rather than with the secretory burden

By expressing μ_s_ we burden the cells with a Sisyphean task, because in absence of λ, no IgM can be produced, a situation that mimics storage of mutant ER client proteins in disease. Inspired by B to plasma cell differentiation, we argued that λ in contrast to μ_s_ would likely provide a poor if any proteostatic insult to the cells, as it is expressed in B lymphocytes when they still are quiescent, and— by definition—minimally stressed. Moreover, ER retention of λ is far less stringent than that of μ_s_ such that λ can be secreted (***Hendershot and Sitia, 2005***). To assess whether the flux of client proteins passing through the ER or rather their accumulation determines the amplitude of UPR signaling, we created clones that co-express the two IgM subunits and selected clones based on stoichiometric differences in λ and μ_s_ expression. As expected, when λ and μ_s_ were co-expressed, antibody assembly took place, which we monitored by immunoblotting of lysates and of culture media separated by reducing (***Figure 3A***) or non-reducing (***Figure 3B***) gel electrophoresis. The clone in which λ was in excess allowed successful assembly into polymeric antibodies that were efficiently secreted. In clones where μ_s_ was in excess, assembly and secretion were less efficient, leading to intracellular accumulation of disulfide-linked species that are characteristic intermediates of the IgM assembly pathway (***Hendershot and Sitia, 2005***). As anticipated, HeLa cells inducibly expressing λ, at similarly high levels as we had reached for μ_s_, did not activate the UPR (***Figure 3C***). Strikingly, the μ_s_ driven UPR was almost fully mitigated as well when λ was co-expressed in excess, while the other clones showed intermediate UPR activation commensurate with the excess of μ_s_ (or lack of λ).

**Figure 3.**
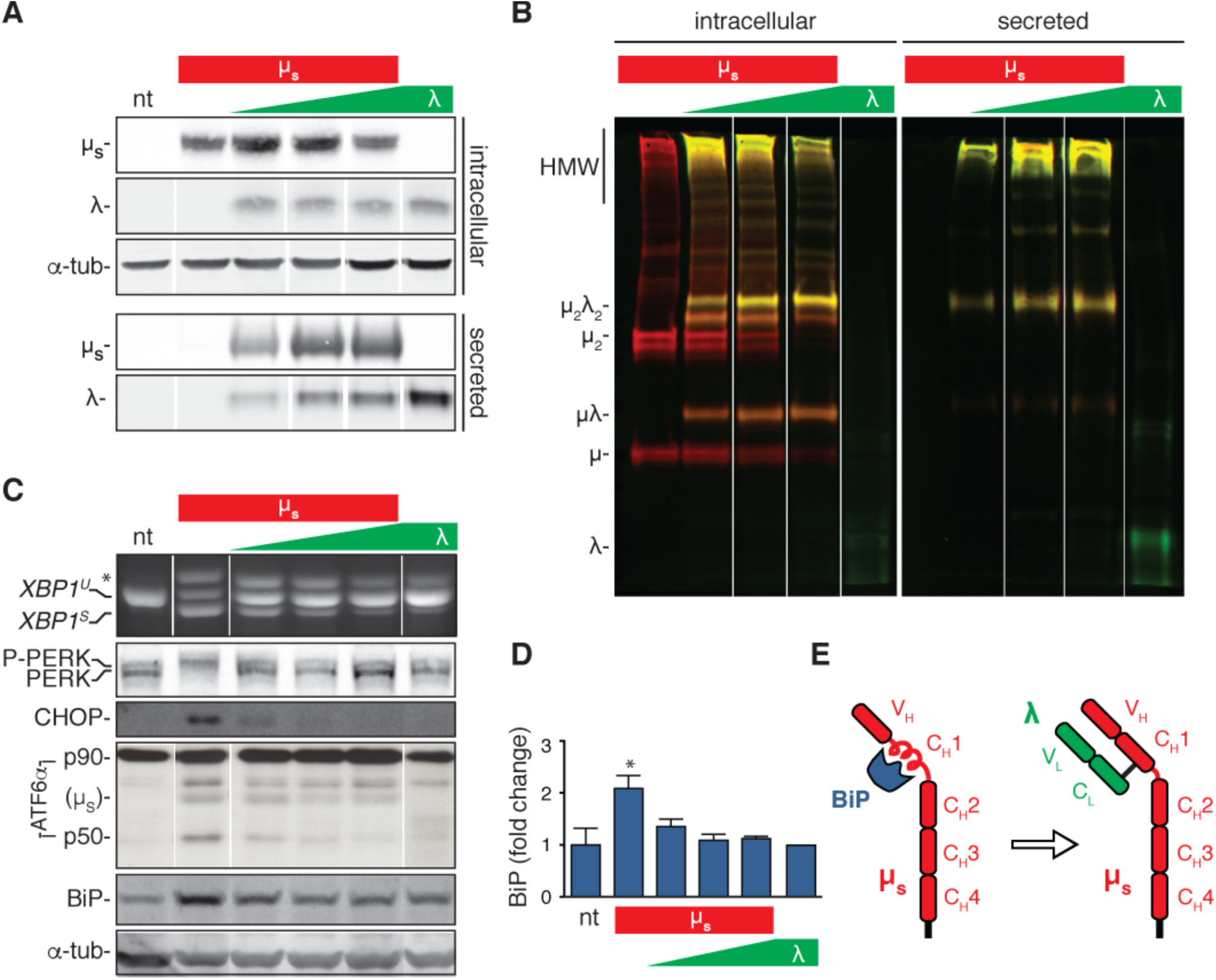
The accumulation of clients in the ER rather than the flux of secretory load drives the UPR. (**A**-**D**) HeLa clones constructed to inducibly express IgL either alone (λ) or in conjunction with μ_s_ (μ_s_ + λ) at different stoichiometries, ranging from μ_s_ being in excess to λ being in excess, all under control of Mif, were induced with 0.5 nM in parallel to the HeLa-μ_s_ cells for 16 hrs to express IgM subunits. Non-treated (nt) HeLa-μ_s_ cells served as a reference. Supernatants (secreted) and cell lysates (intracellular) were separated by reducing (**A**) or non-reducing (**B**) gel electrophoresis. Levels of μ_s_, λ (**A**), CHOP, BiP (**C**) and α-tubulin (**A**,**C**), as well as activation of the UPR pathways (**C**) were assessed as in ***Figure 2A***. (**B**) Immunobloting of μ_s_ (red) and λ (green) reveals disulfide linked assembly intermediates, as indicated: μλ, μ_2_, μ_2_λ_2_, and high molecular weight (HMW) polymeric assemblies of μ_s_ and λ. Simultaneous immunodetection of both IgM subunits resulted in yellow signal. (**D**) Quantitation of BiP levels shown in (**C**) and replicate experiments, mean and s.e.m. are shown, n=3. Statistical significance in a one sample *t*-test of differences in BiP levels is indicated (*p ≤ 0.05). (**E**) Schematic of BiP associating with the CH1 domain of μ_s_ until it is displaced by λ (if available).

Since the levels of ER client proteins (the sum of intracellular λ and μ_s_) were comparable in the different clones, we concluded that the flux of proteins entering the ER is no key determinant for UPR activation, but rather that the nature of the proteostatic insult determines the extent of UPR activation, with λ being a very poor UPR activator, and μ_s_ being a particularly powerful one, provided it does not team up with λ. Of note is that levels of the ER chaperone BiP increased markedly when μ_s_ was expressed alone but far less so or not at all if λ was present (***Figure 3D***). Indeed, it is well-established that in plasma cells BiP binds to μ_s_ until it is displaced by λ (***Bole et al., 1986***; ***Figure 3E***).

### ER homeostatic readjustment entails a transition from acute to chronic UPR signaling

Since the cells that inducibly overexpress μ_s_ showed maximal UPR activation with no major negative impact on cell growth—unlike cells treated with UPR eliciting drugs—we surmised that the μ_s_ expressing cells underwent successful homeostatic readjustment of the ER. To appreciate the timing of the response to the proteostatic insult and the process of ER homeostatic readjustment with a more detailed temporal resolution, we further analyzed by immunoblotting the upregulation of μ_s_, BiP and of another ER chaperone, PDI, with time (***Figure 4A***). As such, we witnessed that levels of BiP only noticeably increased from 12 hrs onwards (and PDI even later), while μ_s_ built up steadily in the first 12 hrs upon induction. Indeed, using recombinant BiP and commercially obtained pure IgM (of which ∼70% of the protein mass is μ_s_) as standards, we assessed the absolute quantities in cell lysates of BiP and μ_s_ upon induction of μ_s_ expression. BiP levels increased ∼12.5-fold. Based on cell counting, protein weight determination of samples and that the molecular weight of BiP is ∼70 kDa, we estimated BiP levels to rise from ∼2 x 10^7^ to ∼2 x 10^8^ copies per cell. Importantly, from the absolute quantitation we deduced that BiP levels were in excess of μ_s_ once the cells had adapted to the proteostatic insult (with μ_s_ reaching an estimated ∼1.5-1.8 x 10^8^ copies per cell). Early upon the onset of μ_s_ expression and before BiP induction was fully underway, however, μ_s_ levels transiently were at a 1:1 stoichiometry with those of BiP, or possibly, μ_s_ levels even slightly exceeded BiP levels (***Figure 4B***).

**Figure 4.**
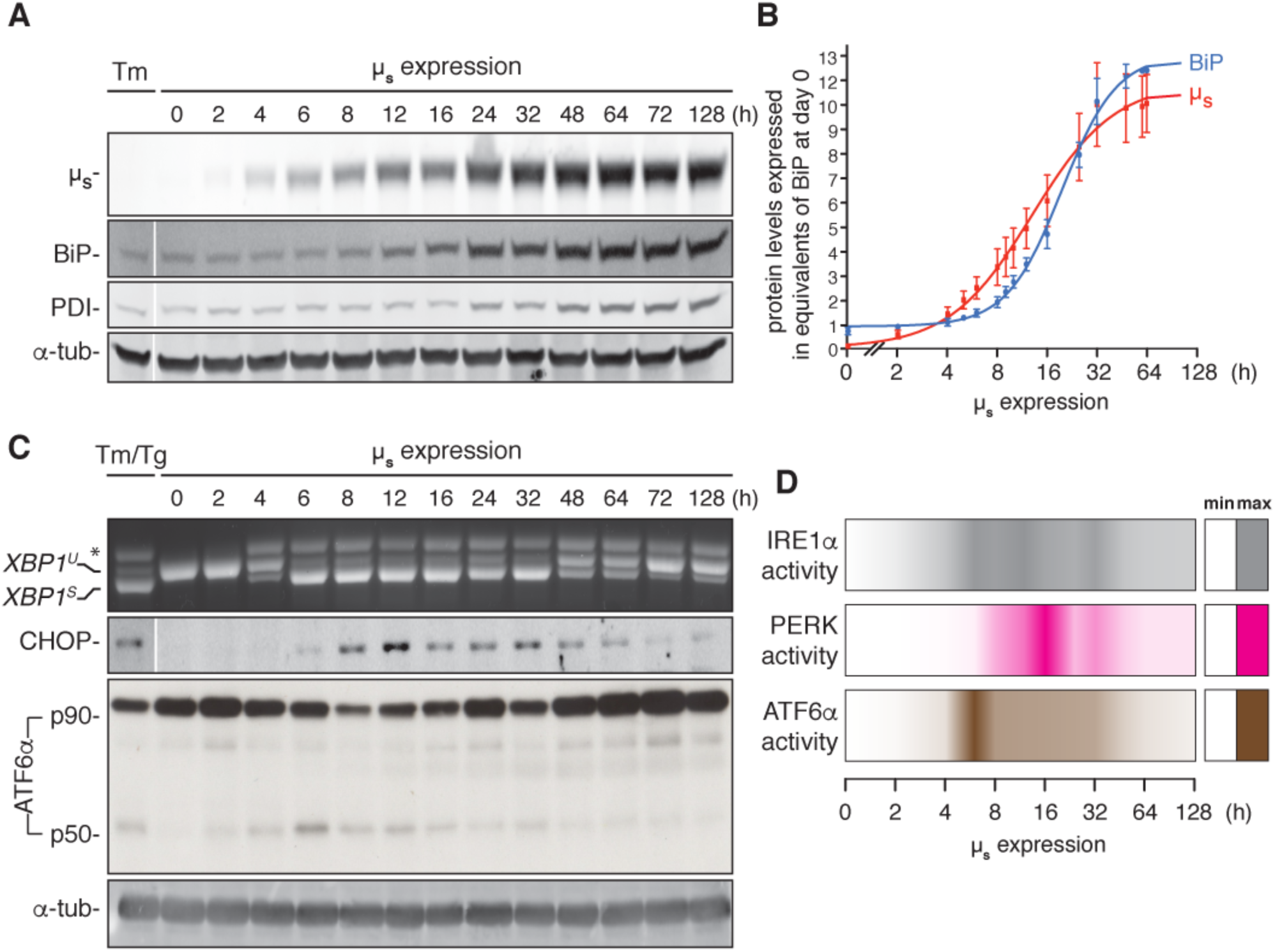
ER homeostatic readjustment entails a transition from acute to chronic UPR signaling. (**A**-**C**) HeLa-μ_s_ cells were induced with 0.5 nM Mif for various times as indicated. Levels of μ_s_, BiP, PDI (**A**) and α-tubulin (**A**,**C**) as well as activation of the UPR pathways (C) were assessed as in ***Figure 2A***. (**B**) Levels of μ_s_ and BiP were quantified using known quantities of BiP and μ_s_ as standards. The X-axis displays time in hrs in a logarithmic scale to show better detail of the early phase of the time course. Experiments were normalized to the level of BiP at *t* = 64 h, which was reproducibly ∼10 fold that at *t* = 0 h. Quantities of BiP and μ_s_ are expressed in equivalents of BiP levels at *t* = 0 h; mean and s.e.m. are shown, n=5. Fitting was done with Prism software to obtain a sigmoidal dose-response curve. Since quantities were not assessed at the same time points in each experiment, some values were inferred from fit curves to obtain the s.e.m. (**D**) Activation of the UPR branches in (**C**) and replicate experiments were quantified (n=2-20, depending on the time point assessed): for IRE1α the percentage of *XBP1* mRNA splicing, for PERK the levels of expression of its downstream effector CHOP, and for ATF6α the levels of ATF6α-p50 (both the latter normalized on the levels of α-tubulin). Mean values of UPR signaling output at each time point were calculated as a percentage of the maximal level, and depicted in a coloring-based heat map ranging from no color (min) to full color (max) displayed on a time bar in logarithmic scale as the X-axis in (**C**).

Following the activation of the UPR over time upon onset of μ_s_ expression, we found that all three UPR branches transiently reached an output that was maximal, i.e. at least on a par with the output that was induced by a conventional drug-elicited UPR (***Figure 4C***). Levels of cleaved ATF6α (p50) reached a maximum at ∼6 hrs upon the onset of μ_s_ expression before it subsided to submaximal levels. Levels of spliced *XBP1* mRNA likewise reached a maximum at the same time but remained maximal for about 24 hrs (with perhaps some oscillations) before they subsided to submaximal levels. Levels of CHOP, which provides a readout for PERK activity, reached a maximum slightly later, at 12-16 hr upon the onset of μ_s_ expression before reaching submaximal levels, as further illustrated in a “heatmap” of UPR activation (***Figure 4D***). Treatment with ER-stress eliciting drugs during the chronic phase when the UPR had submaximal output, readily triggered close to maximal output of all three UPR branches again (***Figure 4—figure supplement 1***), indicating that prolonged μ_s_ expression had not exhausted the signaling capacity of the UPR, and, thus, that homeostatic readjustment of the ER accounted for lowered UPR output levels upon chronic μ_s_ expression. Strikingly, the highest output of the three UPR branches did not coincide when μ_s_ reached maximal expression levels. Instead, UPR signaling reached maxima when the ratio μ_s_/BiP was highest (***Figure 4B,D***). These findings support a scenario in which the UPR signals are commensurate with the extent of μ_s_ sequestering the folding machinery, in particular BiP, rather than its accumulation *per se*.

### The ER expands in response to a proteostatic insult

The increase of BiP levels upon μ_s_ expression indicated that the homeostatic readjustment to accumulating μ_s_ levels entailed expansion of the ER. To directly visualize ER expansion, we targeted a modified version of pea peroxidase (APEX) (***Hung et al., 2014; Lam et al., 2015***) to the ER by use of an N-terminal signal peptide and a C-terminal KDEL. APEX-KDEL was expressed under control of the TetT promoter at such low levels that it did not interfere with the μ_s_-driven UPR (***Figure 5— figure supplement 1***). We then exploited APEX to catalyze polymerization of 3,3’-diaminobenzidine tetrahydrochloride (DAB) upon treatment with H_2_O_2_ to obtain contrast in electron microscopy (EM), and stained as such the ER of cells before or after 1, 3, or 7 days of μ_s_ expression. We determined that the ER lumen before induction of μ_s_ expression occupies 10-12% of the area within the cytoplasm (i.e. excluding the nucleus) in the electron micrographs (***Figure 5***), which by a rough estimate would correspond to (0.10-0.12)^3/2^ ≈ 3-4% of the cytoplasmic volume. Considering that the volume of the cytosol in HeLa cells is estimated to be ∼2x10^3^ μm^3^, i.e. ∼2x10^3^ fl (***Milo, 2013***), we estimated the ER to have a volume of ∼60-80 fl. The ER expanded in the course of 1-3 days upon the onset of μ_s_ expression to cover ∼18% of the cytoplasmic area in the electron micrographs, corresponding to roughly (0.18)^3/2^ ≈ 7-8% of the cytoplasmic volume, but we observed no further ER expansion after three days. Based on these estimates the volume of the ER increased 2-3-fold as a result of μ_s_ expression to ∼120-240 fl per cell.

**Figure 5.**
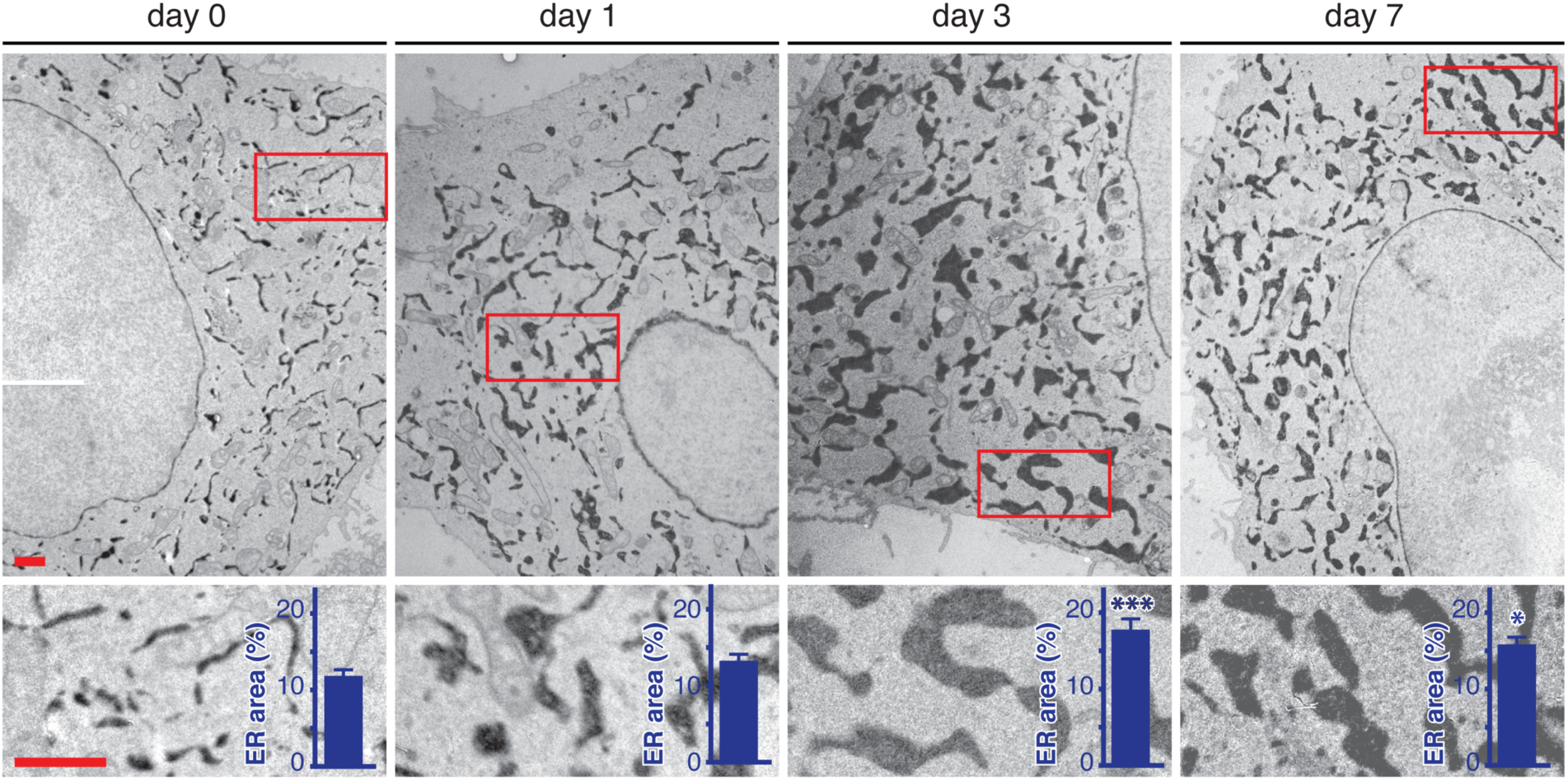
ER expansion in the course of homeostatic readjustment to μ_s_ overexpression. HeLa-μ_s_ derived cells, harboring Dox-inducible APEX-KDEL, were induced with 0.5 nM Mif to express μ_s_ for various days as indicated, and APEX-KDEL expression was induced with 100 nM Dox for 2 days. APEX-KDEL was exploited to obtain DAB precipitates (dark), revealing the extent of the ER in electron micrographs. The percentage of the area within the cytoplasm corresponding to ER was determined and depicted in bar graphs as insets; mean and s.e.m. are shown, n=10. Statistical significance of differences in in the extent of ER occupying cytosolic area in the electron micrographs was tested by ANOVA (*p ≤ 0.05; ***p ≤ 0.001).

### The ER becomes dominated by the chaperone BiP in response to a proteostatic insult

We further corroborated the μ_s_-driven ER expansion by proteomic analysis and identified ∼3000 proteins from lysates of cells before or after 1, 3, or 7 days of μ_s_ expression. We then used a label-free quantitation approach to obtain an approximation of the mass levels of proteins (***Cox et al., 2014***) (***Supplementary File 1***), such that we could estimate what share of the total protein content of the cell the ER would account for. Proteins that we identified at each time point to rank among the 500 most abundant at any of the time points, altogether representing ∼600 proteins, we categorized according to subcellular localization and/or function. We limited detailed analysis to this subset, since the label-free quantitation method is more reliable for abundant proteins (***Cox et al., 2014***), and because this subset already covered >90% of the total protein content in the cell. Changes in protein expression levels were assessed and in parallel by SILAC (***Ong et al., 2002***), which revealed mostly similar trends in differential expression (***Supplementary File 1***), confirming the usefulness of the label-free quantitation approach.

We then calculated for each category the percentage of the total protein content and deduced that the ER accounts for ∼3% before μ_s_ expression in line with the findings by EM. Upon μ_s_ expression the ER content, including μ_s_, expanded to ∼10% within 3 days, but did not expand much further after that (∼12% at day 7) (***Figure 6, upper panels***; ***Supplementary File 1***). The slightly higher estimate of the percentage of the total protein the ER accounts for by proteomics compared to the EM-based estimate of the percentage of the total volume that the ER occupies could indicate that the continuous μ_s_ expression led to increased molecular crowding within the ER. In contrast to the ER, other organelles or cellular machineries were not markedly affected by μ_s_ expression.

**Figure 6.**
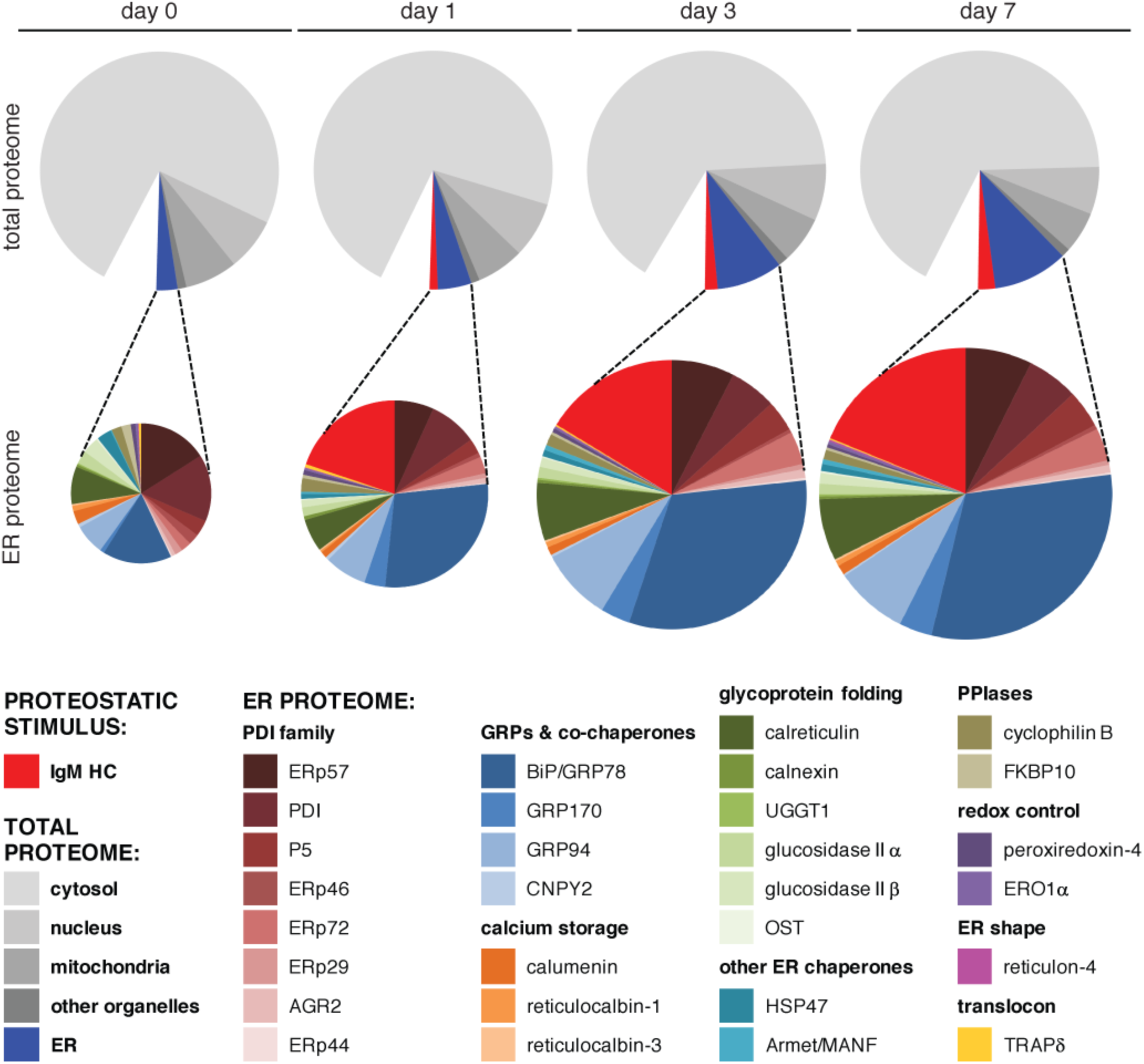
BiP becomes dominant when the ER expands upon μ_s_ overexpression. HeLa-μ_s_ cells were induced with Mif to express μ_s_ for various days as indicated. Proteins at each time point were identified by mass spectrometry and expression levels were approximated by label-free proteomics. The ∼600 most abundant proteins, accounting for over 90% of the total protein content were categorized according to localization in the cell: cytosolic, nuclear, mitochondrial, ER resident, or residing in other organelles by use of Uniprot entries. From the approximated quantities of these proteins the quantities of proteins per organelle were calculated as a percentage of the total proteome and depicted in pie charts (upper panels). For the ER the resident proteins (blue) and μ_s_ (red) are shown separately. Quantities of ER resident proteins including μ_s_ were calculated as a percentage of the ER proteome and depicted in pie charts (lower panels) that are size-proportioned commensurate with the difference in combined quantity of ER and μ_s_ at different days. For details on calculations see: (***Supplementary File 1***). Color-coding of pie charts is as annotated in the legend embedded in the figure.

Levels of ER proteins underwent drastic changes upon μ_s_ expression (***Figure 6, lower panels***; ***Supplementary File 1***). Most notably, levels of BiP increased ∼8-fold (and ∼14-fold according to SILAC measurements; ***Supplementary File 1***) between day 0 and 7, in good agreement with the absolute quantitations we obtained by immunoblotting (***Figure 4B***). In fact, at day 7 BiP became the second most abundant protein in the cells (after γ-actin), amounting to ∼4% of the total proteome. Put together with the absolute quantitation of BiP levels, we estimate the cells on average to harbor 5 x 10^9^ proteins, which fits with earlier assumptions (***Milo, 2013***). Moreover, from these quantitations we estimated BiP levels to be at ∼0.5 mM (or 30-40 mg/ml) before induction and to reach ∼1.5 mM (or 90-120 mg/ml) at day 7, corresponding an overall protein concentration of ∼200-300 mg/ml in the ER. Other ER resident chaperones increased as well, such as GRP94, CRT and some members of the PDI family (most notably ERp72 and P5) albeit to a lesser extent. As a consequence, levels of BiP within the ER increased from ∼15% before onset of μ_s_ expression to 30-40% of the expanded ER proteome. Also levels of GRP170, which acts as a nucleotide exchange factor of BiP (***Behnke et al., 2015***), markedly increased.

Before induction μ_s_ was barely detectable (∼1 ppm) but levels rapidly increased to become ∼10 x 10^3^ ppm, i.e. ∼1% (after 1 day) and ∼2% (from day 3 onward) of the total protein content in the cell (***Figure 6, upper panels***; ***Supplementary File 1***). Strikingly, already after 1 day upon induction of its expression, μ_s_ levels exceeded that of any chaperone in the ER except BiP, which still outmatched μ_s_ 3:2 (***Figure 6, lower panels***; ***Supplementary File 1***). Yet, BiP levels had increased 3-4 fold after 1 day of μ_s_ expression to keep pace with the burden imposed on the ER folding machinery by μ_s_. At later times, the margin of BiP being in excess of μ_s_ increased again, with BiP and μ_s_ together constituting about half of the ER when it had fully expanded. Thus, upon μ_s_ expression in bulk, the ER not only changed quantitatively (i.e. undergoing a ∼3-4-fold expansion), but also qualitatively (i.e. becoming dominated by μ_s_ that is held in check by excess levels of its most devoted chaperone BiP).

### UPR signaling, in particular through ATF6α, is key for ER homeostatic readjustment

Since cell growth and survival were hardly perturbed upon μ_s_ expression, we reasoned that ER homeostasis was successfully readjusted. The HeLa-μ_s_ model therefore permitted to investigate in detail to what extent UPR activation—and which branch in particular—contributes to ER homeostatic readjustment. To that end, we exploited the cells in which IRE1α was deleted and silenced PERK and ATF6α by siRNA (with good efficiency; ***Figure 7—figure supplement 1***), either individually or in combination. Remarkably, ablation of all three UPR branches together *per se* hardly impeded cell growth (***Figure 7A,B***), perhaps because HeLa cells by default handle a low secretory load (***Supplementary File 1***). Concomitant expression of μ_s_, however, fully abolished cell growth, showing that the UPR is essential for ER homeostatic readjustment upon μ_s_ expression. Annexin V staining confirmed that the loss of cell growth was due to apoptosis as μ_s_ expression caused synthetic lethality under conditions when all UPR branches were ablated (***Figure 7C***).

**Figure 7.**
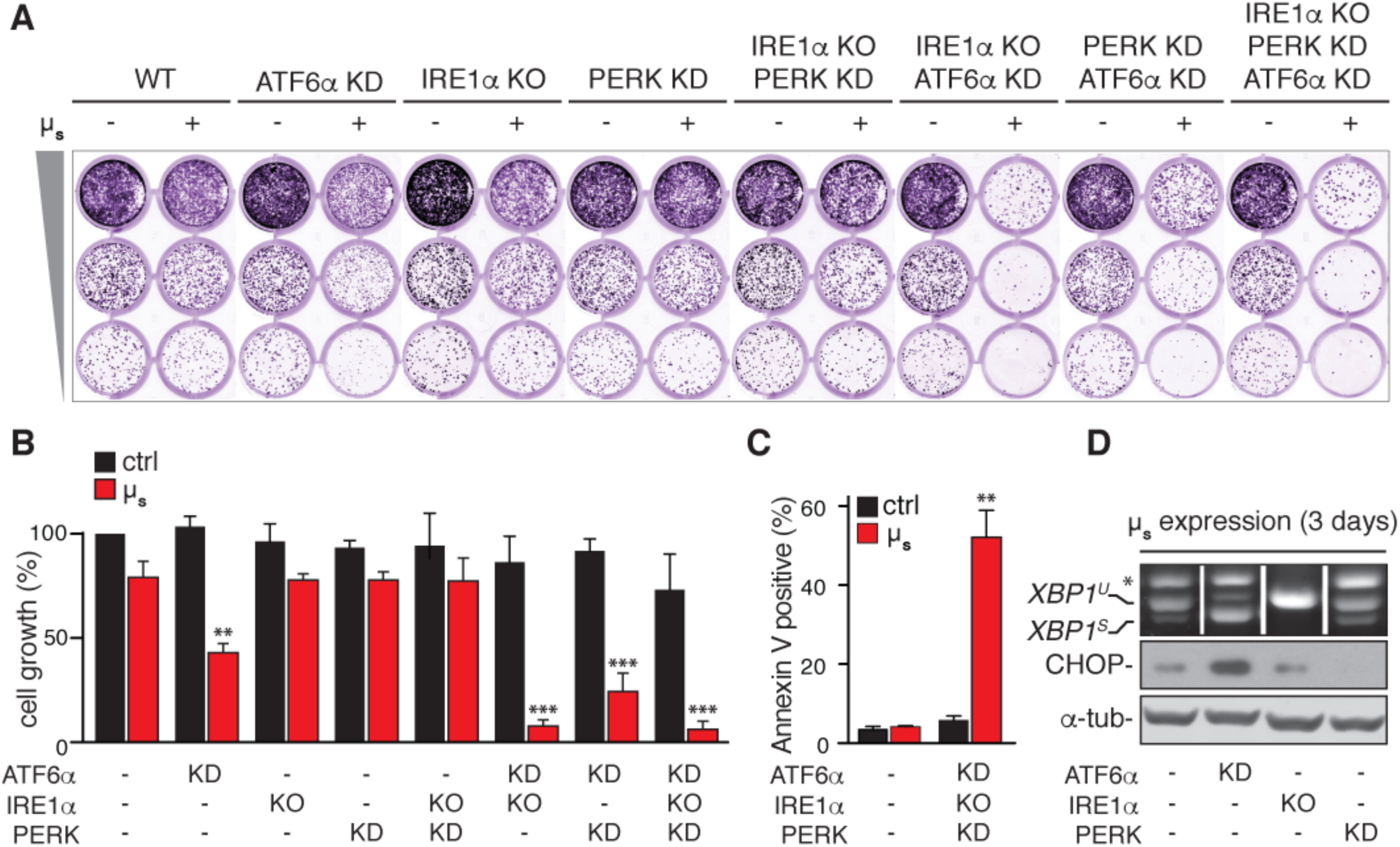
The UPR is essential to sustain ER homeostatic readjustment upon μ_s_ expression. (**A**,**B**) Cell growth was assessed as in ***Figure 1B*** of either wild-type (WT) HeLa-μ_s_ or HeLa-μ_s_ in which UPR transducers were deleted (KO, IRE1α) or silenced (KD, ATF6α and/or PERK) either alone or in combination upon induction of μ_s_ expression with 0.5 nM Mif (+) or not (-), as indicated (**A**), and quantitated as in ***Figure 1C*** (**B**) Mean and s.e.m. are shown in a bar graph; n=2. (**C**,**D**) HeLa-μ_s_ cells in which all three UPR transducers (**C**), or individual UPR transducers (**D**) were ablated as in (**A**) or not (**C**,**D**), upon induction of μ_s_ expression for 3 days were stained with Annexin V (**C**), or activation of UPR pathways was assessed as in ***Figure 2A*** (**D**). (**C**) Percentages of Annexin V positive cells were assessed by cytometric analysis. Mean and s.e.m. are shown in a bar graph, n=2. (**B**,**C**) Statistical significance of differences in growth (**B**) or Annexin V staining (**C**) were tested by ANOVA (**p ≤ 0.01; ***p ≤ 0.001).

Surprisingly, ablation of IRE1α and PERK either alone or in combination could be afforded upon μ_s_ expression, with growth being only marginally affected. In contrast, silencing of ATF6α alone negatively affected growth when μ_s_ was expressed, and additional ablation of IRE1α or PERK almost completely abolished growth. Thus, ATF6α is the key branch of the UPR that sustains ER homeostatic readjustment in the HeLa-μ_s_ model, but the pro-survival role of IRE1α and PERK in letting the cells adjust to μ_s_ expression was unmasked when ATF6α was silenced (***Figure 7A,B***). Still, the lack of ATF6α led to increased ER stress, since both *XBP1* mRNA splicing and CHOP expression were enhanced (***Figure 7D***).

### ER homeostatic readjustment requires expansion of the ER

For conditions that led to a severe loss of cell growth, we obviously could not assess BiP or μ_s_ levels upon induction. Conversely, ablation of either IRE1α or PERK alone or in combination did not diminish BiP induction (***Figure 8A,B***), but the induction of BiP levels was compromised when ATF6α was ablated even though the induction of μ_s_ expression levels was similar (***Figure 8C-E***). Thus, the finding that splicing of *XBP1* mRNA and CHOP induction were elevated when ATF6α was ablated (***Figure 7D***) is in line with the notion that UPR signaling increases commensurate with there being a shortage of BiP compared to μ_s_ (***Figure 4***). Not only did ablation of ATF6α lead to lowered BiP induction, it almost fully abolished ER expansion upon μ_s_ expression (***Figure 8F***). Conversely, μ_s_ driven ER expansion was not compromised upon silencing of IRE1α, PERK or both. Thus, ER expansion, as afforded by UPR signaling—in particular through ATF6α—, is key for ER homeostatic readjustment in adaptation to the proteostatic insult posed on the ER folding machinery by μ_s_.

**Figure 8.**
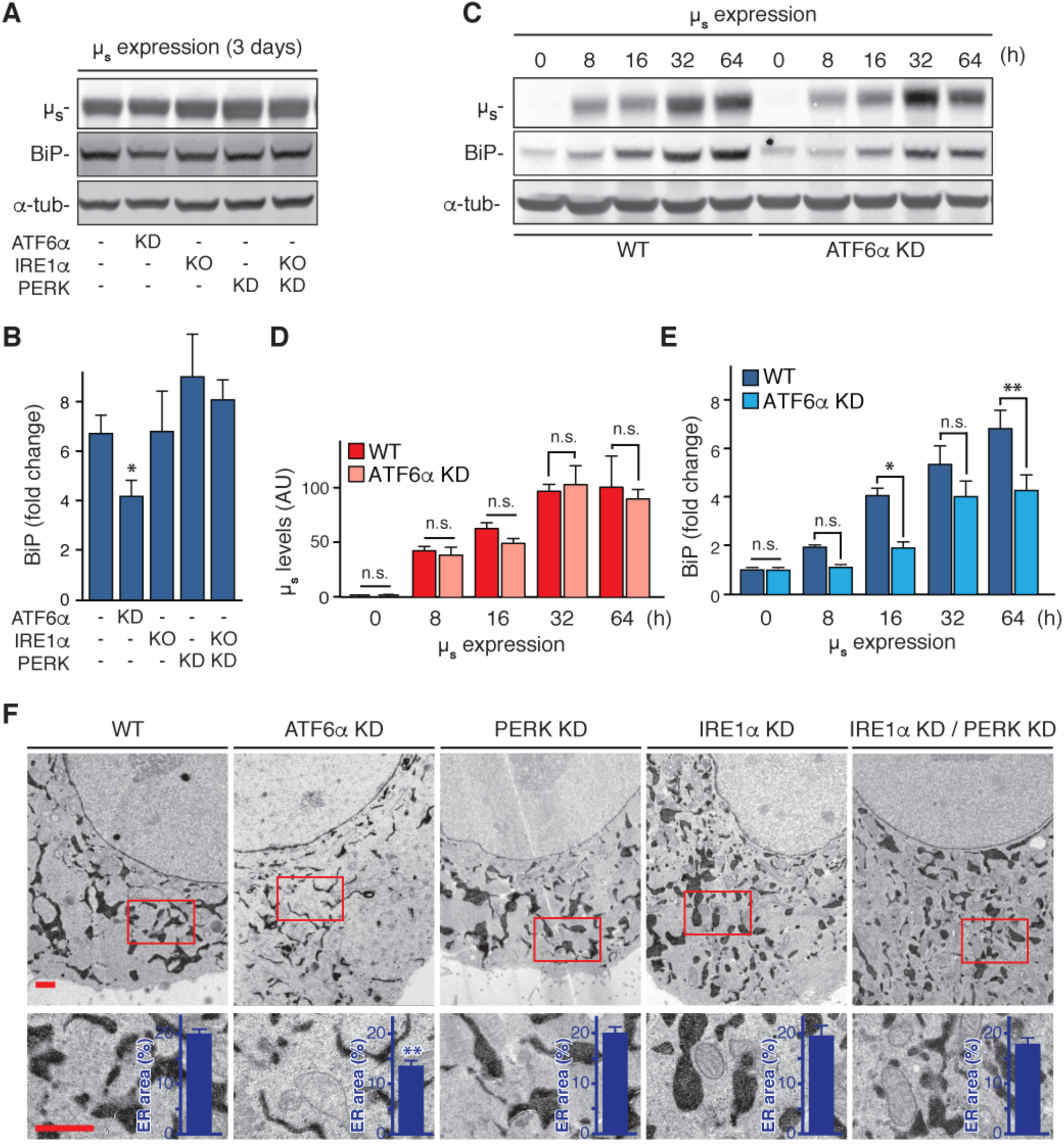
Homeostatic readjustment to cope with μ_s_ expression requires ER expansion. (**A**-**F**). HeLa-μ_s_ cells in which each UPR transducer was ablated alone or in combination, or not (WT), as indicated, were induced with 0.5 nM Mif to express μ_s_ for 3 days (**A**,**B**,**F**) or for the indicated times (**C**-**E**). (**A**,**C**) Levels of μ_s_, BiP and α-tubulin were assessed as in ***Figure 2A***, and BiP (**B**,**E**) and μ_s_, (**D**) levels were quantitated from (**A**), respectively (**C**), and replicate experiments. (**B**,**E**) BiP levels are expressed as fold change compared to untreated cells, as in ***Figure 3D***. (**D**). Levels in WT of μ_s_ at 64 hrs were set at 100. For illustrative purposes this 100 was scaled to levels of BiP in WT at 64 hrs such as to reflect a ratio of μ_s_ to BiP of 2:3, i.e. the average estimate for this ratio based on the absolute quantitations in ***Figure 4B*** (∼7:8), and proteomics ***Figure 6*** (∼1:2). Mean and s.e.m. are shown in bar graphs; n=2-5. (**F**) In cells harboring APEX-KDEL the extent of ER expansion was assessed as in ***Figure 5***. The percentage of the area within the cytoplasm corresponding to ER was determined and depicted in bar graphs as insets. Mean and s.e.m. are shown, n=10-20. (**B**,**D**-**F**) Statistical significance of differences in expression levels (**B**,**D**,**E**), or the extent of ER occupying cytosolic area in the electron micrographs (**F**) were tested by ANOVA (n.s., not significant; *p ≤ 0.05; **p ≤ 0.01).

### ER homeostatic readjustment requires that ER expansion is capped

While ER expansion proved to be essential for the cells to cope with μ_s_ expression, there was no further expansion once a ∼3-4-fold increase of ER content was reached, which implies that at that stage the influx of μ_s_ molecules was matched by countermeasures.

One obvious way through which the stress can be diluted is through cell division, but we reasoned that the cells also employed dedicated mechanisms to achieve a relief of the folding load. Translational attenuation through PERK activation is a means to alleviate the burden on the ER folding machinery by diminishing the input of nascent clients entering the ER lumen (***Harding et al., 1999***). Yet, we ruled out that PERK-driven translational attenuation was a key determinant for ER homeostatic readjustment in the HeLa-μ_s_ model, considering that PERK ablation hardly impeded cell growth upon μ_s_ expression, and considering that there was a transient and only marginal reduction (at the lowest ∼80% of that before induction) in overall protein synthesis only during the first 16 hrs of μ_s_ expression (***Figure 9—figure supplement 1***). RIDD, as afforded by IRE1α, likewise has been proposed to limit ER input by destroying mRNAs that encode ER client proteins (***Hollien and Weissman, 2006; Hollien et al., 2009***). Following the same reasoning, we excluded that RIDD is important for ER homeostatic readjustment upon μ_s_ expression as well, since ablation of IRE1α had little impact on cell growth.

However, μ_s_ is a target of ERAD in plasma cells (***Fagioli and Sitia, 2001***), and we confirmed by radio-labeling-based pulse-chase that μ_s_ was degraded also in the HeLa-μ_s_ cell model in a manner that was almost fully prevented when the proteasomal inhibitor MG132 was present (***Figure 9A,B***). Thus, we ruled out that other degradative pathways (in particular autophagy) play a role in the disposal of μ_s_. Since μ_s_ is glycosylated, it is subject to mannose trimming (***Aebi et al., 2010***), which is a key step in delivering μ_s_ to the dislocation machinery that shuttles μ_s_ to the cytosol for proteasomal degradation (***Fagioli and Sitia, 2001***). Accordingly, the ER mannosidase I inhibitor kifunensine (Kif) stabilized μ_s_ in a similar manner as MG132. Interestingly, while μ_s_ levels built up steadily in the ER with time, ERAD kinetics hardly changed (i.e. the *t*½ of μ_s_ was remarkably constant), which implies that ERAD kept pace with the accumulating load of μ_s_ (***Figure 9C***). Indeed levels of μ_s_ no longer increased (***Figure 4A,B***) soon after the *de novo* synthesis of μ_s_ reached a plateau (***Figure 2E,F***), reflecting that synthesis and degradation were then at equilibrium. The net accumulation of μ_s_ is therefore commensurate with the *t*½ of μ_s_: if the *t*½ would have been shorter, as it is in plasma cells (***Fagioli and Sitia, 2001***), less μ_s_ would accumulate at an equal μ_s_ synthesis rate.

**Figure 9.**
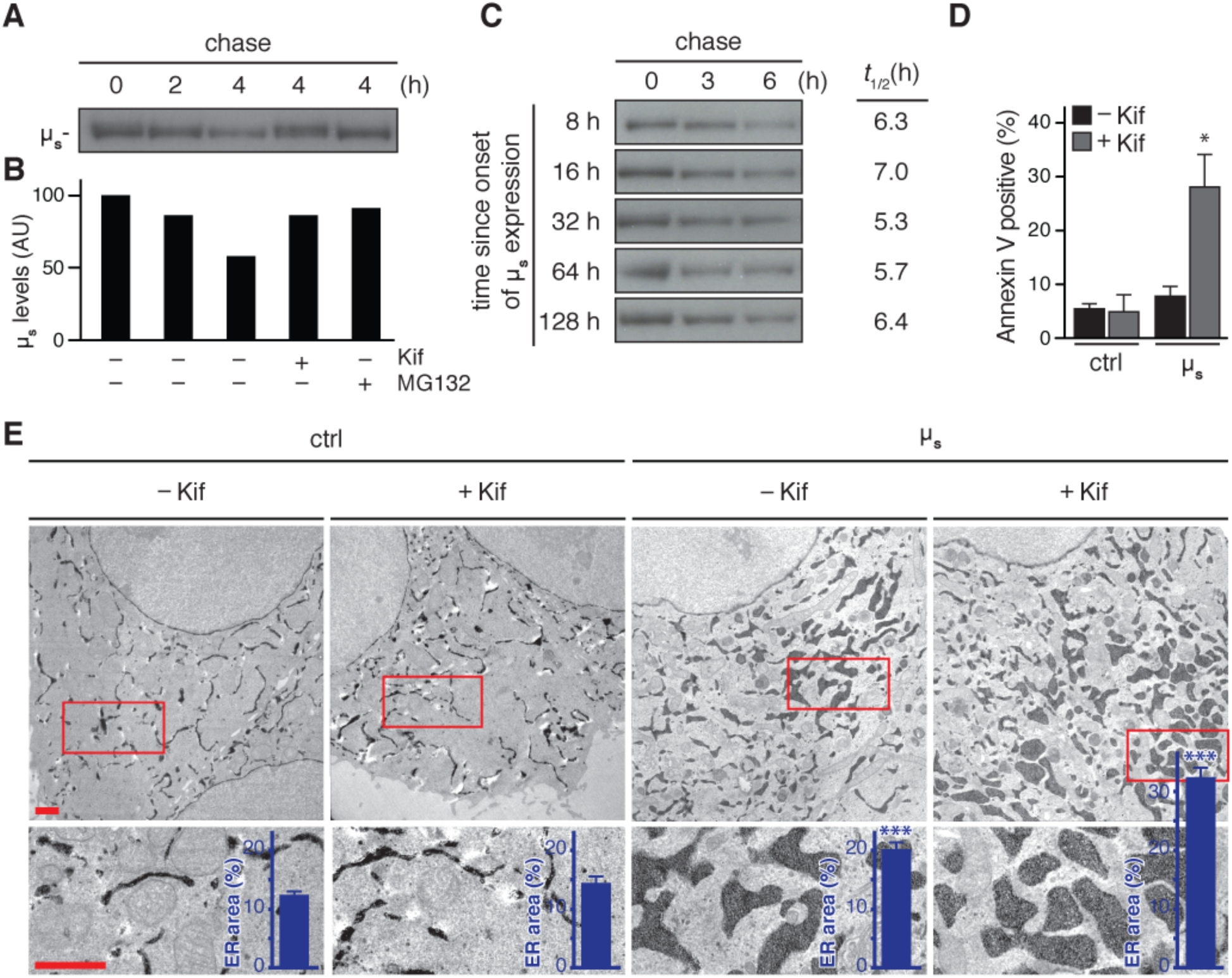
Abrogation of μ_s_ disposal leads to unrestrained ER expansion and cell death. (**A**-**C**) HeLa-μ_s_ cells were pulse labeled for 10 min as in ***Figure 2E*** and chased with excess unlabeled cysteine and methionine for the indicated times at various times after induction with 0.5 nM of μ_s_ expression, in the absence (**A**,**C**) or presence (**A**) of 10 μM MG132 or 30 μM Kif, as indicated. Signals were quantitated as in ***Figure 2E***. (**B**) Quantitation of (**A**) is shown in a bar graph, while quantitations of (**C**) were used to calculate by linear fitting the *t*½ of μ_s_ at various time points after induction of its expression, given on the right. (**D**,**E**) HeLa-μ_s_ cells were not induced (ctrl) or induced with 0.5 nM Mif to express μ_s_ for 3 days in the presence or absence of Kif. (**D**) Percentages of Annexin V positive cells assessed by cytometric analysis, as in ***Figure 7C***. Mean and s.e.m. are shown in a bar graph, n=2-4. (**E**) In cells harboring APEX-KDEL the extent of ER expansion was assessed as in ***Figure 5***. The percentage of the dark area within the cytoplasm corresponding to ER was determined and depicted in bar graphs as insets, mean and s.e.m. are shown, n=10. Statistical significance of differences in Annexin V staining (**D**), or the extent of ER occupying cytosolic area in the electron micrographs (**E**) were tested by ANOVA (*p ≤ 0.05; ***p ≤ 0.001).

Kif treatment did not affect cell viability in HeLa-μ_s_ cells in absence of μ_s_ expression, but caused apoptosis when μ_s_ expression was induced, indicating that ERAD is key for homeostatic readjustment of the ER when under duress of accumulating μ_s_ (***Figure 9D***). Accordingly, ablation of either HRD1 or SEL1L, two ERAD-related proteins shown previously to target μ_s_ for proteasomal degradation (***Cattaneo et al., 2008***), was tolerated in the HeLa-μ_s_ cells, but not upon μ_s_ expression (***Lari et al, manuscript in preparation***).

Not surprisingly, a block of ERAD by Kif entailed further expansion of the ER to an extent that—as judged by electron microscopy—is reminiscent of that in professional secretory cells (***Figure 9E***). The area of ER staining within the cytoplasm reached 30-35%, which on a rough estimate would account for (0.3-0.35)^3/2^ ≈ 17-20% of the cytoplasmic volume, implying that the ER had expanded ∼6-7-fold. In contrast, the extent of the ER was hardly affected by Kif in absence of μ_s_ expression, which further indicates that the default burden on the ER folding machinery in HeLa cells is low. We concluded that curtailing the μ_s_ load by ERAD is essential for the cells to cope with μ_s_ expression. We moreover surmised that the UPR might play a role in capping ER expansion by invoking pro-apoptotic pathways in response to unrestrained ER expansion.

### Unbridled ER expansion causes IRE1α and PERK to relinquish their pro-survival role

Since we found that ATF6α was key for sustaining ER expansion to accommodate the accumulating μ_s_ load, and, since ERAD turned out to be a crucial countermeasure, it was not surprising that ablating ATF6α and ERAD at the same time resulted in further loss of cell growth than when either ATF6α was silenced or ERAD inhibited alone (***Figure 10A***,***B***). Unexpectedly, ablation of either IRE1α or PERK, and, in particular, of both IRE1α and PERK led to a partial “rescue”; i.e. inhibition of ERAD caused less reduction in growth upon μ_s_ expression when the cells lacked IRE1α and PERK. We therefore surmised that under conditions when ERAD is dysfunctional, IRE1α and PERK are no longer predominantly pro-survival, but instead contribute to a loss in cell growth. Thus, when μ_s_ cannot be disposed of and ER expansion is unrestrained, IRE1α and PERK apparently commit to a pro-apoptotic role. Successful homeostatic readjustment of the ER entailed that signaling through IRE1α and PERK subsided to a submaximal level after an acute but transient phase of maximal signaling (***Figure 4A***,***C***). In contrast, ablation of ERAD caused IRE1α and PERK to chronically signal at maximal levels when μ_s_ was expressed (***Figure 10C***). In line with the vast ER expansion we witnessed under those conditions (***Figure 9E***), BiP levels augmented a further ∼2-fold when ERAD was inhibited but to no avail, since μ_s_ levels underwent an even greater, ∼3-fold, rise, likely resulting in a relative shortage of BiP (***Figure 10D***), which suggests once more that maximal activation of the UPR transducers coincides with BiP running short.

**Figure 10.**
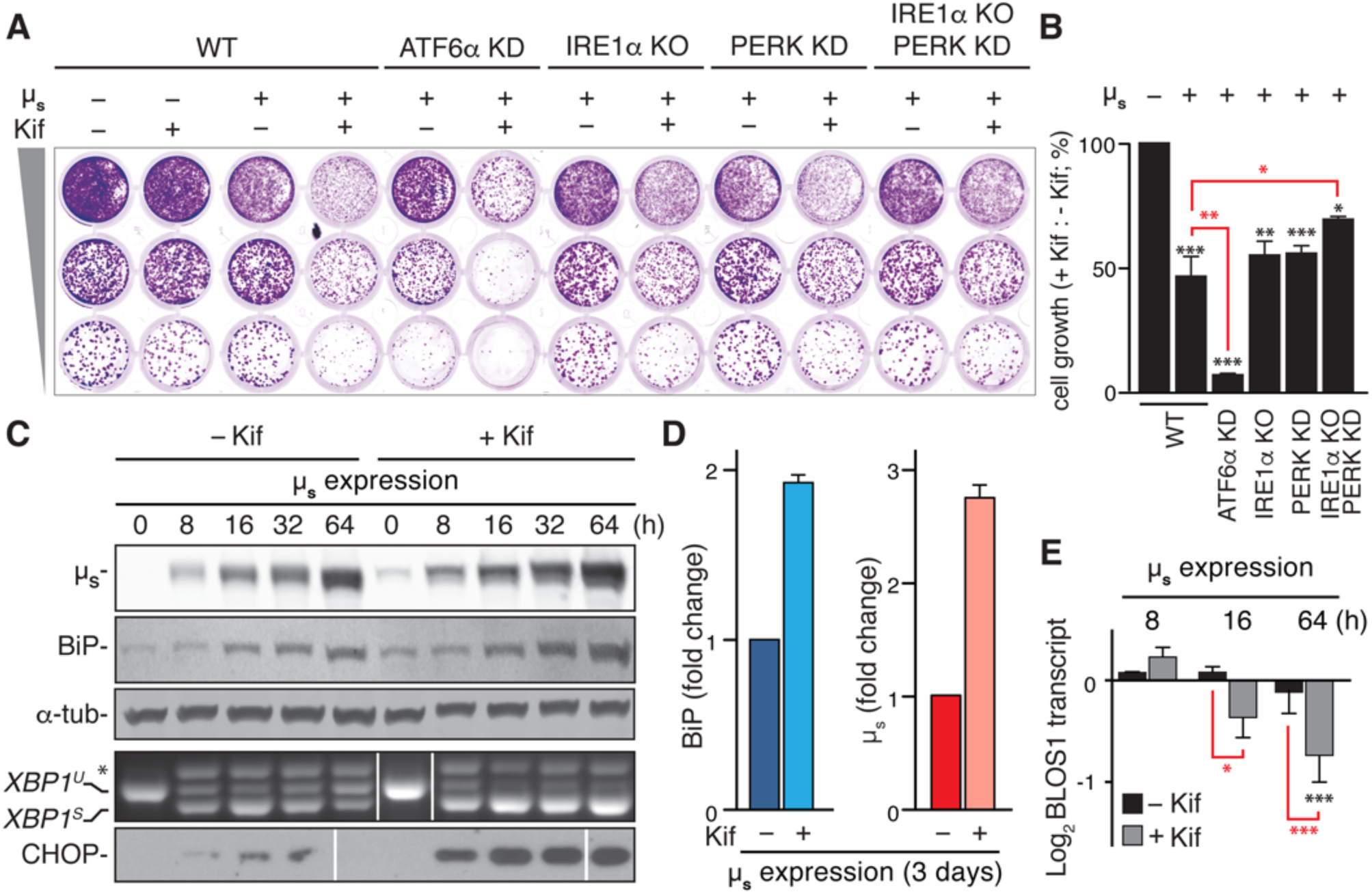
Upon unrestrained ER expansion IRE1α and PERK commit to a pro-apoptotic role. (**A**) Growth of HeLa-μ_s_ cells in which each UPR transducer was ablated alone or in combination, or not (WT) were induced with 0.5 nM Mif to express μ_s_ or not in the absence or presence of 30 μM Kif, as indicated, was assessed as in ***Figure 7A***. (**B**) Growth in (**A**) and replicate experiments was quantitated by assessing crystal violet staining. For each condition the ratio of growth upon inhibition of ERAD with Kif versus no ERAD inhibition was determined. The ratio for WT cells that did not express μ_s_ was set at a 100%. Mean and s.e.m. are shown in a bar graph; n=3. (**C**-**E**) HeLa-μ_s_ cells were induced for various times as indicated (**C**,**E**) or for 3 days (**D**) in the absence or presence of 30 μM Kif. (**C**) Levels of μ_s_, BiP, and α-tubulin as well as activation of the IRE1α and PERK branches of the UPR were assessed as in ***Figure 2A***. (**D**) Levels of BiP and μ_s_ were assessed by absolute quantitation as in ***Figure 4B*** and depicted in bar graphs as in ***Figure 7D***, such that the μ_s_ levels in the absence of Kif were scaled to BiP levels at a ratio of 2:3. Levels in the presence of Kif are expressed as a fold change compared to levels in the absence of Kif; mean and s.e.m. are shown; n=2. (**E**) Levels of *BLOS1* mRNA relative to untreated cells were determined by real time PCR. Mean and s.e.m. are shown in a bar graph; n=2-5. (**B**,**D**,**E**) Statistical significance of differences in the ratio of growth in the presence or absence of Kif between all conditions compared to WT cells that either express μ_s_ (red) or not (black) (**B**), of differences in protein levels comparing – Kif and + Kif conditions (**D**), and of differences in *BLOS1* mRNA levels compared to untreated cells (black) or compared between Kif treated or non treated μ_s_ expressing cells at the same time point (red) (**E**) were tested by ANOVA (**B**,**E**), or a Student’s *t*-test (**D**) (*p ≤ 0.05; **p ≤ 0.01; ***p ≤ 0.001).

Reportedly, unmitigated PERK activation leads to cell death as continuous attenuation of protein synthesis—not surprisingly—is detrimental for cell growth and survival (***Lin et al., 2009***). Moreover, constitutively high levels of CHOP are implicated in provoking cell death (***Zinszner et al., 1998***). How rampant IRE1α activation ties in with pro-apoptotic signaling, however, is still debated even though several mechanisms have been proposed, including RIDD activity (***Han et al., 2009***). Indeed, when μ_s_ was expressed in bulk, it transiently provoked decay of *BLOS1* mRNA, which is indicative of IRE1α having committed to RIDD activity (***Hollien et al., 2009***). Yet, RIDD activity was unabated upon chronic μ_s_ expression only when also ERAD was ablated, and, therefore, ER homeostatic readjustment failed (***Figure 10E***).

### IRE1α commits to pro-apoptotic activity through RIDD

To confirm in a direct manner that RIDD activity accounts for IRE1α-provoked loss of cell growth, we exploited that the IRE1α KO cells that had been reconstituted with IRE1α-GFP allow to titrate levels of IRE1α-GFP by Dox. Indeed, commensurate with its overexpression, IRE1α-GFP provoked a loss of cell growth (***Figure 11A,B***). The overexpression of IRE1α-GFP led to its auto-activation, causing *XBP1* mRNA splicing to reach maximal levels (***Figure 11C,D***) even in the absence of ER stress, as has been reported before (***Han et al., 2009; Li et al., 2010***). The loss of cell growth was not exerted through *XBP1* mRNA splicing, however, since silencing of XBP1 actually aggravated the loss of cell growth (***Figure 11A***,***B***). Yet, treatment with 4μ8C, which inhibits IRE1α’s endonuclease function (***Cross et al., 2012***; ***Figure 11C,D***), almost fully restored cell viability (***Figure 11A***,***B***), annulling the effects of IRE1α-GFP overexpression.

**Figure 11.**
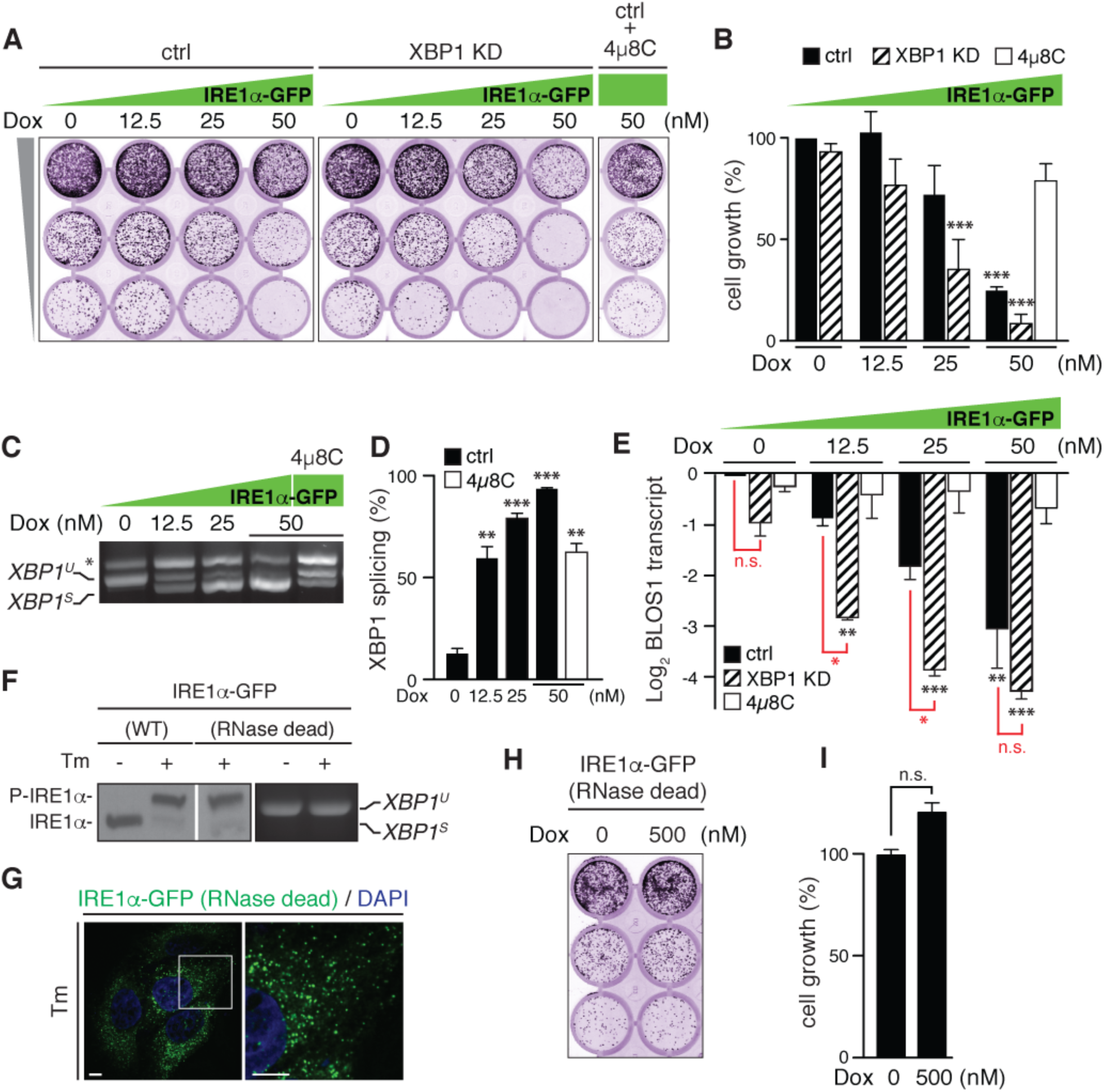
*IRE1*α *commits to a pro-apoptotic role through RIDD activity when levels of unspliced XBP1 mRNA are exhausted*. (**A**) Growth assay as in ***Figure 1B*** of HeLa cells in which endogenous IRE1α was ablated but reconstituted with IRE1α-GFP, as in ***Figure 2H***, that were induced with increasing concentrations of Dox, as indicated, to incrementally overexpress IRE1α-GFP. Expression of *XBP1* mRNA was silenced (XBP1 KD) or not (ctrl) (**A**-**C**) Endonuclease activity of IRE1α was inhibited with 4μ8C where indicated. (**B**) Growth in (**A**) and replicate experiments was quantitated as in ***Figure 1C***. Mean and s.e.m. are shown in a bar graph; n=4. (**C**-**E)** RNA was extracted from cells treated for 48 hrs under the same conditions as in (**A**) to assess *BLOS1* mRNA levels (**E**), as in ***Figure 10E***, and *XBP1* mRNA splicing (except when it was silenced), as in ***Figure 1A*** (**C**). The percentage of the total *XBP1* mRNA that was spliced was quantitated as described (***Shang and Lehrman, 2014***) (**D**). Means and s.e.m. are shown in bar graphs; n=2. (**B**,**D**,**E**) Statistical significance of differences in growth (**B**), or the percentage of *XBP1* mRNA splicing (**D)**, or *BLOS1* mRNA levels (**E**) compared to untreated cells (black), or between cells that were silenced for XBP1 or not at otherwise identical conditions (red) were tested by ANOVA (*p ≤ 0.05; **p ≤ 0.01; ***p ≤ 0.001). (**F**-**I**) HeLa cells in which endogenous IRE1α was ablated but reconstituted with Dox inducible IRE1α-GFP (WT), as in ***Figure 2H***, (**F**), or IRE1α-GFP (RNase dead) (**F**-**I**) were non-treated (**F**,**H**,**I**) or treated with Dox to express the transgenes at comparable, near-endogenous levels (0 nM, WT; 5 nM, RNase dead) (**F**); 25 nM to allow visualization of IRE1α-GFP (RNase dead) by fluorescence microscopy, as in ***Figure 2H***, (**G**); or 500 nM (**H**,**I**), and treated for 4 hrs with 5 μg/ml Tm, as indicated, (**F**,**G**) or not (**F**,**H**,**I**). Lysates were separated by Phos-tag gel electrophoresis and phosphorylation of IRE1α-GFP was detected by a shift to a lower mobility form, P-IRE1α, in the immunoblot (**F**, left). Splicing of *XBP1* mRNA was analyzed as in ***Figure 1A*** (**F**, right). (**G**) Nuclei were stained with DAPI. Scale bars represent 10 μm. The right panel is a 3-fold magnification of the micrograph shown on the left. (**H**) Growth assay as in ***Figure 1B*** of the IRE1α-GFP (RNase dead) reconstituted cells in which the transgene was highly overexpressed with 500 nM Dox or not. (**I**) Quantitation of the growth assay in (**H**) assessed as in ***Figure 1C***. Mean and s.e.m. are shown in a bar graph; n=2. There is no statistical significance (n.s.) in a Student’s *t*-test of differences in growth between conditions.

Loss of cell growth instead correlated well with the extent of RIDD activity (as measured by decay of *BLOS1* mRNA) that was caused by the overexpression of IRE1α-GFP (***Figure 11E***), and RIDD activity and loss of growth were further enhanced when XBP1 was silenced (***Figure 11A,B,E***). These findings confirm that RIDD activity is triggered by unconstrained activation of IRE1α and support that RIDD activity of IRE1α is sufficient for provoking apoptosis, as reported (***Han et al., 2009; Oslowski et al., 2012; Lerner et al., 2012; Ghosh et al., 2014***). Importantly, *XBP1* mRNA splicing activity and RIDD activity are not mutually exclusive. Instead, IRE1α seems to switch from pro-survival to pro-apoptotic activity, simply when it is in short supply of its preferred substrate unspliced *XBP1* mRNA whether due to silencing or because most *XBP1* mRNA is already spliced.

Earlier reports have shown that when IRE1α is activated and phosphorylated it can engage various pro-apoptotic pathways (***Tabas and Ron, 2011***), for instance by recruiting the adaptor molecule TRAF2 that then can couple IRE1α to ASK1. ASK1 in turn can activate JNK, a transcription factor that heralds cell death (***Urano et al., 2000***). Alternatively or in parallel, TRAF2 reportedly couples IRE1α phosphorylation with caspase-7 and caspase-12 activation, which likewise promote apoptosis (***Yoneda et al., 2001***). We then reconstituted the IRE1α KO cells with an endonuclease activity deficient (N906A; ***Han et al., 2009***) mutant of IRE1α-GFP, which upon Tm treatment was phosphorylated despite its lack of splicing activity (***Figure 11F***) and which clustered into foci (***Figure 11G***) to a similar extent as wild-type IRE1α Α ***Figure 2H***), but did not provoke loss of cell growth even upon substantial overexpression (***Figure 11H,I***). These findings further support that IRE1α commits to a pro-apoptotic role exclusively by means of RIDD.

## DISCUSSION

By avoiding the use of ER stress-eliciting drugs that have cytotoxic side effects, we have created a cell model that allowed us to reevaluate several aspects of how cells respond to an accumulating load of client protein in the ER. First, our results highlight that the UPR predominantly acts as a pro-survival pathway. Namely, by confronting cells with a proteostatic stimulus (i.e. bulk μ_s_ expression) that these cells could well cope with, we confirmed that the UPR is key for their capacity to do so. Yet, the finding that μ_s_ expression readily causes cells to undergo apoptosis in the absence of these UPR transducers, implies that other, as yet unknown ER stress-induced mechanisms can be invoked to mediate cell death, which may well play such a role also when the UPR is functional. For instance, hyperoxidizing conditions in the lumen of the ER activate NADPH oxidases, leading to efflux of radical oxygen species to the cytosol that may cause activation of pro-apoptotic pathways (***Tabas and Ron, 2011***).

Second, our data confirm that the UPR drives ER expansion, which—in agreement with earlier reports on non-secretory cells (***Wu et al., 2007; Yamamoto et al., 2007***)—mainly relied on the ATF6α branch in the HeLa-μ_s_ model. Interestingly, ER expansion is not necessarily provoked by bulk synthesis of client proteins *per se*, but rather by their accumulation, as co-expression of sufficient quantities of IgL led to bulk secretion of IgM, but failed to trigger a strong UPR, and, thus, did not result in augmented BiP production (which served as a proxy for ER expansion). Our findings indicate moreover that the ER folding machinery seems to be underused in HeLa cells. Apparently, mobilization of the default ER folding machinery is sufficient for a task that can be productively executed, such as the folding and assembly of IgM from subunits that are expressed in bulk but in the correct stoichiometry. In most cells the basal size of the ER and its folding capacity indeed may exceed the need for folding assistance from the default secretory burden. In line with such a notion, a substantial fraction of BiP is kept in reserve in an inactive, AMPylated form under non-ER stress conditions (***Preissler et al., 2015***).

ERAD genes are UPR targets, and, accordingly, ERAD activity increased in the HeLa-μ_s_ model such that it kept pace with the increasing μ_s_ influx in the sense that ERAD sustained a remarkably constant *t*½ of μ_s_ regardless of its expression levels. It has been argued that kinetics of ERAD are tuned such that a delay in their disposal offers incompletely folded or assembled ER clients a grace period to mature after all (***Ellgaard and Helenius, 2001***). The inadvertent consequence of ERAD’s leniency is that ER clients accumulate if they never will reach maturity, such as orphan μ_s_, causing ER expansion. It recently has become clear that autophagy can serve to curtail ER expansion (***Pengo et al., 2013; Khaminets et al., 2015; Fumagalli et al., 2016***), but in our model disposal of μ_s_ appeared to be exclusively afforded through ERAD, and, even when ERAD was inhibited, we found no indications that autophagy contributed as a backup to hold excess μ_s_ expression in check. This finding implies that specific signals are required for activation of ER-phagy, which are not engendered when μ_s_ accumulates in the ER of HeLa cells.

Third, our results highlight that a shortage of BiP in matching the accumulating load of μ_s_ correlated with maximal UPR activation, rather than accumulation of client protein in the ER *per se*. Indeed, the transitioning of UPR signaling to chronic submaximal levels coincided with the ER becoming dominated by BiP. Our findings are in line with the idea—which was first coined by David Ron and colleagues (***Bertolotti et al., 2000***)—that sequestering of BiP by ER clients (such as μ_s_) is key for activating the UPR, since activation of the main three UPR sensors, IRE1α, PERK (***Bertolotti et al., 2000***) and ATF6α (***Shen et al., 2002***) goes hand in hand with BiP dissociation from their luminal ER stress sensing domains, while overexpression of BiP dampens UPR activation (***Bertolotti et al., 2000***). Accordingly, the PiZ mutant of α1-antitrypsin despite its accumulation in the ER does not sequester BiP and fails to trigger the UPR (***Graham et al., 1990***).

Based on the crystal structure of yeast Ire1’s lumenal domain (***Credle et al., 2006***) Peter Walter and colleagues proposed that Ire1 activates through direct binding of unfolded proteins, for which evidence has been obtained in the meantime (***Gardner and Walter, 2011***). While we cannot exclude that ATF6α, IRE1α and/or PERK directly interact with μ_s_, it unlikely would be the main, let alone exclusive, mechanism of UPR activation, as it is difficult to envision how the interaction between μ_s_ and the UPR transducers would diminish during chronic μ_s_-driven ER stress when μ_s_ is more abundant in the ER than any other protein except for BiP. A recent study, moreover, supports that sequestering of BiP by μ_s_ indeed is sufficient to activate IRE1α and PERK (***Carrara et al., 2015***).

Our choice to employ μ_s_ as a proteostatic insult to challenge the ER was in fact motivated also by the reasoning that if there ever would be an ER client protein to be excellent in sequestering BiP, it would be μ_s_ (***Haas and Wabl, 1983; Bole et al., 1986***). Furthermore, it is well-conceivable that even the slightest leakiness of unpaired μ_s_ from the ER has been evolutionarily selected against. Since antigen recognition likely is compromised if μ_s_ is released from the cell when unaccompanied by λ, it potentially would lead to ill-fated off-target effects. Thus, BiP binding to μ_s_ must be highly stringent out of immunological necessity (***Anelli and van Anken, 2013***), which correlates with the fact that mutations in μ_s_ that lead to lowered affinity for BiP are associated with disorders of the immune system, such as lymphoproliferative heavy chain disease or myeloma (***Hendershot et al., 1987; Anelli and Sitia, 2010***).

Apparently to ascertain stringent ER retention of the accumulating load of μ_s_, the pool of BiP and the ER at large indeed expanded to an impressive extent, in particular when ERAD was inhibited. In fact, the excessive ER expansion in the HeLa-μ_s_ model upon preventing ERAD of μ_s_ is reminiscent of that in plasma cells. In the course of B to plasma cell differentiation all cellular machineries change to anticipate bulk IgM production (***van Anken et al., 2003***). Conversely, the purely μ_s_-driven stimulus we investigated in this study resulted almost exclusively in ER expansion. We found only meager indications even for expansion of the remainder of the secretory machinery (***Supplementary File 1***). The stringent retention of μ_s_ in the ER indeed obviates the need for reinforcing the secretory pathway downstream of the ER. Accordingly, homeostatic regulation of the Golgi and beyond appears to be governed foremost by dedicated signaling mechanisms other than the UPR (***Luini and Parashuraman, 2016***).

Our findings furthermore provide insight into two important aspects of ER stress signaling that are relevant for disease. First, various genetic disorders stem from mutations in ER client proteins that lead to their misfolding and accumulation in the ER. Based on our results, cells that suffer from such conditions will likely display submaximal UPR signaling levels similar to what we found for chronic μ_s_ expression, as prolonged maximal UPR signaling correlates with invoking pro-apoptotic pathways. Thus, in cells or tissues that express a mutant or orphan protein that accumulates in the ER, submaximal UPR signaling is not a sign of “mild” ER stress, but rather reflects that the stress these cells suffer from has been sub-lethal, such that they have adapted to the stress that is chronic, although the stress likely still predisposes cells to succumb.

Second, our data emphasize that in the HeLa-μ_s_ model IRE1α commits to a pro-apoptotic role through RIDD activity rather than through processing of *XBP1* mRNA or through IRE1α being in a phosphorylated state, which may play role in other cells instead (***Tabas and Ron 2011***). Importantly, little if any RIDD activity accompanied successful homeostatic readjustment of the ER. IRE1α activity *per se* contributed marginally to successful ER homeostatic readjustment. Thus, in the HeLa-μ_s_ model RIDD activity unlikely serves any protective role, neither by decreasing the burden on the ER folding machinery by depleting mRNA pools encoding for ER client proteins (***Hollien et al., 2009***) nor by degrading specific target mRNAs, such as has been shown for death receptor 5 (***Lu et al., 2014***). Unrestrained RNase activity of IRE1α against RNA targets other than *XBP1* mRNA may in itself cause a plethora of problems to the cell that ultimately lead to activation of pro-apoptotic pathways. Yet, RIDD activity reportedly also invokes cell death in a selective manner through cleavage of various microRNAs (***Upton et al., 2012***). Particularly intriguing is RIDD-controled decay of miR17, as it leads to the stabilization of the mRNA encoding TXNIP (***Lerner et al., 2012***). Since TXNIP is upregulated at the same time through PERK and its downstream effector ATF5 (***Oslowski et al., 2012***), PERK and IRE1α pathways converge in promoting apoptosis via TXNIP-driven triggering of the NLRP3 inflammosome and ensuing caspase activation. Ablation of this signaling relay in fact prevents ER stress-induced apoptosis in pancreatic β cells (***Oslowski et al., 2012; Lerner et al., 2012***).

Altogether, a dichotomy emerges between IRE1α’s splicing activity being pro-survival and its RIDD activity being pro-apoptotic. Strategies that would selectively inhibit or promote either of IRE1α’s activities therefore would hold therapeutic promise. Indeed, it has been proposed that IRE1α transitions from pro-survival *XBP1* mRNA splicing activity to pro-apoptotic RIDD activity by virtue of a switch in its enzymatic activity (***Han et al., 2009; Tam et al., 2014; Maurel et al., 2014***). Our data indicate however that once IRE1α commits to RIDD activity, *XBP1* mRNA splicing activity is still unabated, which argues against any such switch. Instead, we argue that IRE1α’s endonuclease activity is unleashed against other RNA substrates simply because IRE1α’s preferred substrate, unspliced *XBP1* mRNA, is running low. Such a scenario would be attractive also to explain how the extent of ER expansion is capped depending on the maximal levels of UPR specific transcription factors, in particular ATF6α-p50 and XBP1 (***Yamamoto et al., 2007***), that can be mobilized to drive ER expansion. If *XBP1* mRNA splicing levels are already maximal but the UPR driven ER expansion still cannot keep pace with the increasing demand on the ER folding machinery, IRE1α activity is directed to other RNA substrates and thus automatically commits to RIDD to invoke pro-apoptotic pathways.

Along these lines, levels of ATF6α, IRE1α and *XBP1* mRNA could serve as key determinants in defining the maximal extent of ER expansion and sensitivity to ER stress of a given cell. Indeed, transcript levels of ATF6α (http://www.gtexportal.org/home/gene/atf6) and of IRE1α (http://www.gtexportal.org/home/gene/ern1) differ between cell types by approximately one order, while XBP1 (http://www.gtexportal.org/home/gene/xbp1) differs by more than two orders of magnitude. The ratio of *XBP1* mRNA to IRE1α levels is much higher in secretory tissues than in non-secretory cells. As such, quasi-inexhaustible supplies of spliceable *XBP1* mRNA increase the threshold for IRE1α to commit to RIDD, thereby protecting secretory cells better against ER expansion-induced apoptosis, while affording through the XBP1 protein further reinforcement of the ER folding capacity. Pro-apoptotic RIDD activity may be even further avoided by fueling IRE1α’s endonuclease with a decoy RNA in case no unspliced *XBP1* mRNA is left. For instance, μ_s_ mRNA in plasma cells becomes a RIDD target once XBP1 is ablated (***Benhamron et al., 2014***). Following the same reasoning, non-secretory cells would be highly susceptible to RIDD-driven cell death, since the stocks of unspliced *XBP1* mRNA will more readily run short. If these cells—uncharacteristically—expand their ER too much, RIDD provides a means for self-censorship. An attractive therapeutic avenue against RIDD-driven cell death therefore would be to employ a competitive inhibitor or decoy substrate for which IRE1α has an affinity that is outmatched only by its affinity for unspliced *XBP1* mRNA.

## MATERIAL AND METHODS

### Cell culture

HeLa S3 cells and all derivate lines were cultured in DMEM (Gibco-Life Technologies) containing glutamax (1 mM), 5% Tet-System approved Fetal Bovine Serum (FBS, ClonTech), 100 U/ml penicillin and 100 µg/ml streptomycin. Expression of transgenes was induced with 0.5 nM Mif unless indicated otherwise, and/or Dox at various concentrations as indicated. I.29μ^+^ lymphomas were used as model B lymphocytes (***Alberini et al., 1987***), and cultured in suspension in RPMI (Gibco-Life Technologies) supplemented with 10% LPS free FBS (Hyclone), glutamax (1 mM), penicillin (100 U/ml), and streptomycin (100 μg/ml), sodium pyruvate (1 mM) and β-mercaptoethanol (50 μM). I.29μ^+^ cells were induced to differentiate with 20 μg/ml LPS (Sigma).

### Generation of HeLa derived cell lines that inducibly express transgenes

The pSwitch cassette (GeneSwitch system; Invitrogen), placed into a lentiviral vector, as described (***Sirin and Park, 2003***)—a kind gift from Dr. Frank Parker—, was used to render cells Mif responsive, as the pSwitch cassette encodes a hybrid nuclear receptor that is activated by Mif to drive expression of genes under the control of the GAL4 promoter. Similarly, a reverse tetracycline-dependent transactivator (rtTA) cassette (***Zhou et al., 2006***) under control of a bidirectional promoter with ΔLNGFR in the reverse orientation for selection purposes (***Amendola et al., 2005***) on a lentiviral vector was employed to render cells Tet (and thus Dox) responsive. A pGene5b (GeneSwitch system; Invitrogen) encoding lentiviral construct (***Sirin and Park, 2003***)—another kind gift from Dr. Frank Parker—was used as backbone to place either Igμ_s_ or Igλ under control of the GAL4 promoter. The coding sequences for NP-specific murine μ_s_ and λ were derived from plasmids described elsewhere (***Sitia et al., 1987; Fagioli and Sitia, 2001***). pENTR223.1 #100061599 bearing murine IRE1α cDNA was obtained from Thermo Fisher Scientific. GFP was placed in frame at the same position in the juxtamembrane cytosolic linker portion of IRE1α as described (***Li et al., 2010***). The IRE1α-GFP cassette was placed under control of a TetTight (TetT) responsive element (Clontech) into a lentiviral vector. This construct was mutagenized to harbor the N906A substitution in the endonuclease domain of IRE1α to obtain an RNase dead version of IRE1α-GFP (***Upton et al., 2012***). The APEX2 coding sequence (***Lam et al., 2015***), derived from the #49385 plasmid (Addgene), and modified to contain in frame extensions encoding the vitronectin signal peptide at the N-terminus and the tetrapeptide KDEL at the C-terminus, was placed under control of a TetT promoter into the same lentiviral vector as employed for IRE1α-GFP. Standard techniques were used for construction, transformation and purification of plasmids. Transgene cassettes were genomically integrated in a subsequent manner into HeLa S3 cells by lentiviral delivery, essentially as described (***Amendola et al., 2005***). Cells with genomic integrations of transgenes were cloned by limiting dilution to yield the cell lines used in this study as summarized in (***Supplementary File 2A***).

### Cell growth and apoptosis assays

To assess growth, cells were counted with a Burker chamber, and seeded in 24-multiwell plates in 1:5 serial dilutions (5000, 1000, and 200 cells per well) to grow for 7 days. Culture media and pharmacological agents were refreshed every 2-3 days. Cells were fixed in methanol-acetone (1:1) for 10 min, stained with 0.5% crystal violet in 20% methanol for 10 min and washed with distilled water. Dried plates were densitometrically scanned at a resolution of 50-100 µm with a Typhoon FLA-9000 reader (GE Healthcare), employing the 647 nm laser and the photomultiplier 1000. Intensity of crystal violet staining was analyzed with ImageJ for quantitation of growth. Typically, wells originally seeded with 1000 cells were used for comparison of signal intensity between conditions. An empty well on the same plate served for background subtraction. As a measure for apoptosis the percentage of Annexin V positive cells was recorded by use of a standard detection assay (APC-Annexin V, BD Biosciences) on a Canto cytometer (BD Biosciences) following the manufacturers’ instructions.

### *XBP1* mRNA splicing assay and RIDD assay by real-time PCR of *BLOS1* mRNA

Total RNA was extracted from cells using the UPzol RNA lysis reagent (Biotechrabbit) followed by a standard protocol provided by the manufacturer for assessing *XBP1* mRNA splicing levels, or by using the DirectZol kit with on-column DNAse I treatment (Zymo Research) for BLOS1 RT-PCR. From samples cDNA was obtained and amplified by PCR. Oligos used to amplify cDNA corresponding to *XBP1* mRNA have been described (***Calfon et al., 2002***). PCR products were resolved on agarose gels and images were acquired with the Typhoon FLA-9000 reader. The percentage of spliced *XBP1* mRNA was calculated as described (***Shang and Lehrman, 2014***). Relative levels of *BLOS1* mRNA were determined by use of a ViiA7 ThermalCycler with pre-mixed primers and probe (Hs01041241_g1; both Applied Biosystems), according to the standard protocol; 18S RNA (Hs99999901_s1; Applied Biosystems) served as a reference for normalization. Quantitation was done as described (***Livak and Schmittgen, 2011*)**.

### Protein analysis

Protein extraction, sample preparation, electrophoresis on 10% or 4-12% Bis/Tris precast polyacrylamide gels (Novex, Life Technologies), and transfer of proteins onto nitrocellulose (GE Healthcare Life Science) for immunoblot analysis with antibodies listed in ***Supplementary File 2B*** were performed using standard techniques with the following exceptions: for analysis of CHOP and ATF6α cells total lysates were used to ensure nuclear pools of these proteins were solubilized; for analysis of disulfide-linked IgM assembly intermediates, reducing agents were omitted from the lysis buffer and free sulfhydryl groups were alkylated with N-ethyl-maleimide (NEM); for analysis of ATF6α 1.5 mm thick 8% polyacrylamide gels, prepared from a 30% polyacrylamide-bis 29:1 solution (Biorad), were used as described (***Maiuolo et al., 2011***); for analysis of IRE1α phosphorylation cells were lysed with 50 mM TrisHCl (pH 7.4), 150 mM NaCl, 60 mM octylglucoside (all from Sigma) and proteins were separated on 4% acrylamide gels containing 25 μM Phos-tag^TM^ reagent (WAKO), prepared according to the manufacturers’ instructions. Protein transfer onto nitrocellulose was confirmed by reversible Ponceau staining. Protein quantitation of samples was performed using a bicinchoninic acid (Sigma).

Detection of fluorescent antibody signals on blots was performed by scanning with the Typhoon FLA-9000 reader, except for IRE1α and ATF6α, which were detected on films using HRP-conjugated secondary antibodies and ECL (Amersham). Signal intensities were analyzed with Image J. For absolute quantitation of BiP and μ_s_ protein levels standard μ_s_ and BiP curves were obtained. To that end we used purified mouse myeloma derived IgM (#02-6800, Invitrogen), of which we estimated 70% of the protein mass to be μ_s_, and purified recombinant hamster BiP, which safe for two conservative changes, Y313F and A649S, is identical in sequence to human BiP and therefore, in all likelihood, is recognized equally well by the goat anti-BiP antibody (C-20; Santa Cruz). Recombinant BiP was expressed and purified as described (***Marcinowski et al., 2011***) from a pPROEX expression construct that was a kind gift from Dr. Johannes Buchner. To avoid potential crossreaction of secondary antibodies against μ_s_ we used an anti-IgM antibody that was itself Alexa-546-conjugated to allow fluorescent detection.

### Fluorescence microscopy

Samples were prepared for immunofluorescence as described (***Sannino et al., 2014***), and sample containing coverslips were mounted on glass slides with Mowiol (Sigma). Light microscopic images were acquired with an UltraView spinning disc confocal microscope operated by Volocity software (PerkinElmer). Images were processed with Photoshop (Adobe). Antibodies used were: Alexa-546 anti-mouse IgM (μ) 1:1000 (Life Technologies); rabbit anti-calreticulin 1:200 (Sigma), secondary Alexa-488-anti-rabbit 1:300 (Life Technologies). Nuclei were stained with DAPI (Sigma).

### Ablation of UPR pathways and ERAD, and pharmacological activation of the UPR

By CRISPR/Cas9 endogenous IRE1α was deleted in HeLa S3 or derivate cells (***Supplementary File 2A***), and clones were obtained by limiting dilution and verified by their lack of *XBP1* mRNA splicing upon Tm treatment. Gene silencing of ATF6α, IRE1α, PERK, and XBP1 was performed with pooled ON-TARGET plus siRNA (Dharmacon) (***Supplementary File 2C***) according to the manufacturer’s instructions. Cells were seeded for experiments the day after transfection with the siRNA pools. The inhibitor of IRE1α’s endonuclease activity, 4μ8C (a kind gift from Dr. David Ron), and the α-mannosidase inhibitor Kif (Sigma), which abrogates ERAD of μ_s_, were used at concentrations of 10 μM, and 30 μM respectively. The UPR was pharmacologically elicited with either Tg (Sigma) at a concentration of 300 nM, or Tm (Sigma) at a concentration of 5 μg/ml or lower, as indicated.

### Radiolabeling and pulse-chase

Cells were starved for 10 min in standard medium, but lacking cysteine and methionine, and containing 1% dialyzed FBS, prior to 10 min pulse labeling with Express [^35^S] Protein labeling mix (Perkin Elmer) containing 40 μCi ^35^S-methionine and ^35^S-cysteine per 10^6^ cells. Cells were either harvested immediately, or washed twice in ice-cold HBSS (Gibco-Life technologies), and grown in OptiMEM (Gibco-Life technologies) for various chase times. After a wash in ice-cold PBS, cells were lysed in RIPA buffer, containing NEM and protease inhibitors for 10 min on ice, as described (***Fagioli and Sitia, 2001***).

Lysates were cleared for 10 min at 13,000 rpm at 4°C. Before immunoprecipitation, cell lysates were pre-cleared for 1 hr with 30 µl of FCS-Sepharose (GE healthcare) and µs was immunoprecipitated for 16 hrs using a rabbit anti-mouse IgM (H) antibody (#61-6800, Life Technologies). Immunoprecipitates were collected on protein G-Agarose beads (Invitrogen), washed twice in 10 mM Tris (pH 7.4), 150 mM NaCl, 0.5% NP-40 and once in 5 mM Tris-HCl (pH 7.5) before gel electrophoresis. Gels were transferred and dried onto a 3MM filter paper, and exposed to a LE storage phosphor screen (GE Health Care). Signals were acquired on the Typhoon FLA-9000 with a phosphorimaging filter. Densitometric analysis of signals was performed with ImageJ.

### Electron microscopic determination of ER size

Cells harboring APEX2-KDEL were fixed in 1% glutaraldehyde, 0.1 M sodium cacodylate (pH 7.4) for 30 min, and incubated for 20 min with 0.3 mg/ml DAB, and for 20 min with 0.03% H2O2 (to activate APEX) in 0.1 M sodium cacodylate (pH 7.4), all at room temperature. Samples were rinsed in 0.1 M sodium cacodylate (pH 7.4), and post-fixed with 1.5% potassium ferrocyanide, 1% OsO4, sodium cacodylate (pH 7.4) for 1 hr on ice. After en bloc staining with 0.5% uranyl acetate overnight at 4°C in the dark, samples were dehydrated with increasing concentrations of ethanol, embedded in EPON and cured in an oven at 60°C for 48 h. Ultrathin sections (70 – 90 nm) were obtained using an ultramicrotome (UC7, Leica microsystem, Vienna, Austria), collected, stained once more with uranyl acetate and Sato’s lead solutions, and visualized in a transmission electron microscope (Leo 912AB, Carl Zeiss, Oberkochen, Germany). Digital micrographs were taken with a 2Kx2K bottom mounted slow-scan camera (ProScan, Lagerlechfeld, Germany) controlled by the EsivisionPro 3.2 software (Soft Imaging System, Münster, Germany). Images of randomly selected cells (10 for each condition) were acquired at a nominal magnification of 1840x. Using ImageJ software, cytoplasmic regions were selected manually and ER profiles, thanks to the electron-dense DAB precipitate, were segmented by means of an automatic macro based on threshold intensity, yielding the percentage of the area within the cytoplasm that was ER. For simplicity we used the formula: volume (%) = (area (%))^3/2^—which would be fully accurate only if the ER were cuboidal, but which we considered a good approximation nonetheless—to obtain a rough estimate of the ER volume.

### Proteomics analysis

SILAC heavy and light collected serum-free cell media were mixed proportionally to a 1:1 cell number ratio, lysed with 8 M Urea 10 mM Tris-HCl buffer, and proteins were quantified by Bradford. About 5 μg of mixed proteins for each sample were reduced by TCEP, alkylated by chloroacetamide, digested by Lys-C and trypsin (***Kulak et al., 2014***), and peptides were desalted on StageTip C18 (***Rappsilber et al., 2003***). Samples were analyzed in duplo on a LC–ESI–MS-MS quadrupole Orbitrap QExactive-HF mass spectrometer (Thermo Fisher Scientific). Peptides separation was achieved on a linear gradient from 93% solvent A (2% ACN, 0.1% formic acid) to 60% solvent B (80% acetonitrile, 0.1% formic acid) over 110 min and from 60 to 100% solvent B in 8 min at a constant flow rate of 0.25

μl/min on UHPLC Easy-nLC 1000 (Thermo Scientific) connected to a 23-cm fused-silica emitter of 75 μm inner diameter (New Objective, Inc. Woburn, MA, USA), packed in-house with ReproSil-Pur C18-AQ 1.9 μm beads (Dr Maisch Gmbh, Ammerbuch, Germany) using a high-pressure bomb loader (Proxeon, Odense, Denmark). MS data were acquired using a data-dependent top 20 method for HCD fragmentation. Survey full scan MS spectra (300–1650 Th) were acquired in the Orbitrap with 60000 resolution, AGC target 3^e6^, IT 20 ms. For HCD spectra, resolution was set to 15000 at *m*/*z* 200, AGC target 1^e5^, IT 80 ms; NCE 28% and isolation width 2.0 *m*/*z*. For identification and quantitation Raw data were processed with MaxQuant version 1.5.2.8 searching against the database uniprot_cp_human_2015_03 + sequences of μ_s_ and the Mif responsive hybrid nuclear receptor Switch setting labeling Arg10 and Lys8, trypsin specificity and up to two missed cleavages. Cysteine carbamidomethyl was used as fixed modification, methionine oxidation and protein *N*-terminal acetylation as variable modifications. Mass deviation for MS-MS peaks was set at 20 ppm. The peptides and protein false discovery rates (FDR) were set to 0.01; the minimal length required for a peptide was six amino acids; a minimum of two peptides and at least one unique peptide were required for high-confidence protein identification.

The lists of identified proteins were filtered to eliminate known contaminants and reverse hits. Normalized H/L ratios were analyzed via Perseus (version 1.5.0.6). Statistical analysis was performed using the Significance B outlier test where statistical significance based on magnitude fold-change was established at p < 0.05. To look for proteins that changed over time we considered Intensity L and Intensity H normalized by the correspondent Summed Intensity and the statistical ANOVA test analysis was done using Perseus (ver. 1.5.0.6) with p < 0.01. All proteomic data as raw files, total proteins, and peptides identified with relative intensities and search parameters have been loaded into Peptide Atlas repository (ftp://PASS01009:PJ3566i@ftp.peptideatlas.org/). To obtain approximations for absolute protein quantities we followed a MaxLFQ label-free quantification strategy, as described (***Cox et al., 2014***). We then assessed the relative abundance of total protein per subcellular compartment, i.e. cytosol, nucleus, mitochondrion, ER, and other organelles, as well as μ_s_, as well as the relative abundance of protein species, including or excluding μ_s_, within the ER as detailed in the legend of ***Supplementary File 1***.

## ACKNOWLEDGEMENTS

We thank Drs Bernhard Gendtner, Frank Park, David Ron, and Johannes Buchner for sharing plasmids or reagents. We thank Dr Caterina Valetti for her advice on immunoblotting of ATF6α, Dr Céline Schaeffer for her advice on the use of Phos-tag, Dr John Christiansson for help with RNA silencing, and Dr Andrew Basset for gRNA design. All members of the ALEMBIC imaging facility and the Anna Rubartelli, Paola Panina, Sitia, Bachi and Van Anken labs are acknowledged for stimulating discussions and advice. We thank Drs Maurizio Molinari, Alison Forrester, John Christiansson, Tomás Aragón, and Luca Rampoldi for proofreading of the manuscript.

### COMPETING INTERESTS

The authors declare no competing interests.

## Figure supplements

**Figure 2—figure supplement 1.**
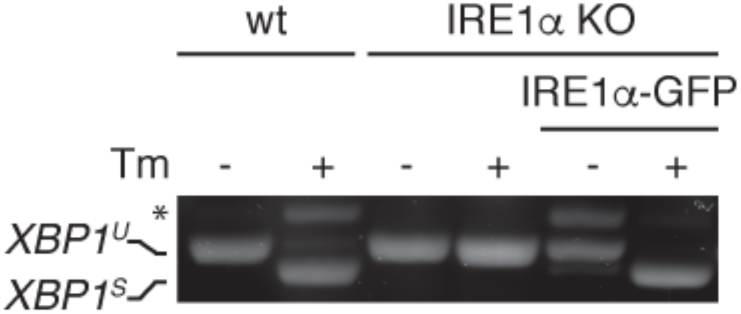
Functional reconstitution of IRE1α-KO cells with IRE1α-GFP. Wild-type (WT) or IRE1α knock out (KO) HeLa cells whether reconstituted with IRE1α-GFP or not were treated (+) or not (-) with 5μg/ml Tm for 4 hrs. *XBP1* mRNA splicing was assessed as in ***Figure 1A***. Functional reconstitution did not require Dox induction of IRE1α-GFP, since “leaky” expression was sufficient.

**Figure 5—figure supplement 1.**
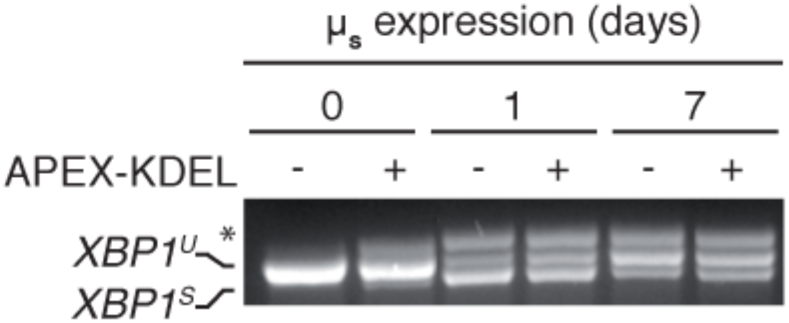
APEX-KDEL expression hardly interferes with μ_s_-driven UPR. HeLa-μ_s_ cells expressing APEX-KDEL induced at low levels with 100 nM Dox for 2 days (+) or not (-) were induced with 0.5 nM Mif to express μ_s_ for the indicated times (days), and *XBP1* mRNA splicing was assessed as in ***Figure 1A***.

**Figure 4—figure supplement 1.**
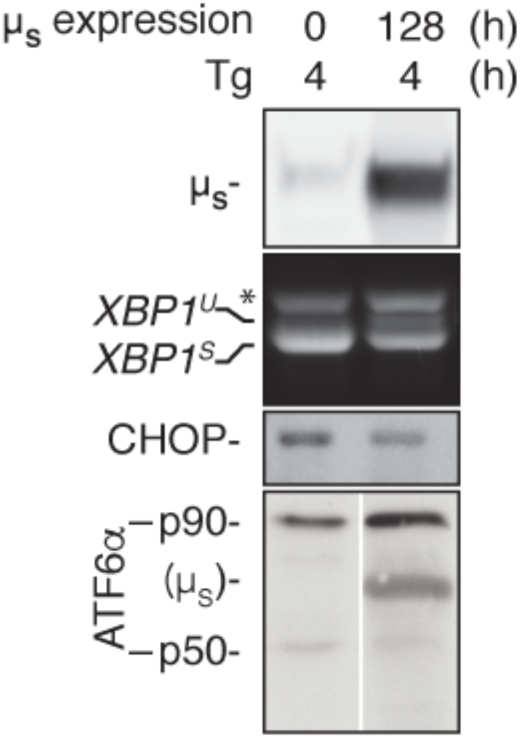
UPR pathways are not exhausted upon chronic μ_s_ overexpression. HeLa-μ_s_ cells were induced for 128 hrs to express μ_s_ or not prior to treatment with 300 nM Tg for 1.5 hrs. Levels of μ_s_ and activation of the UPR pathways were assessed as in ***Figure 2A***.

**Figure 7—figure supplement 1.**
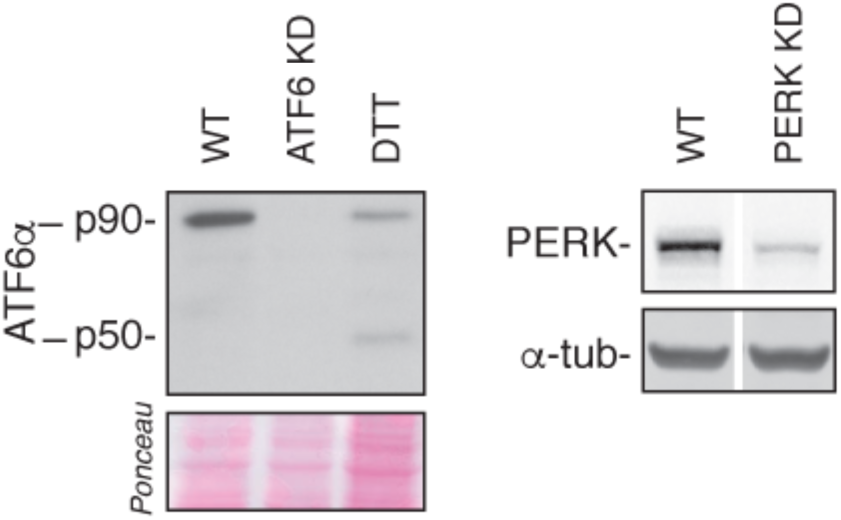
Efficiency of ATF6α and PERK silencing. HeLa-μ_s_ cells were transfected with siRNAs (KD) targeting ATF6α or PERK or not (WT). Levels of ATF6α and PERK were assessed as in ***Figure 2A***; 1 mM DTT treatment for 1 hr was used as a positive reference for ATF6α cleavage.

**Figure 9—figure supplement 1.**
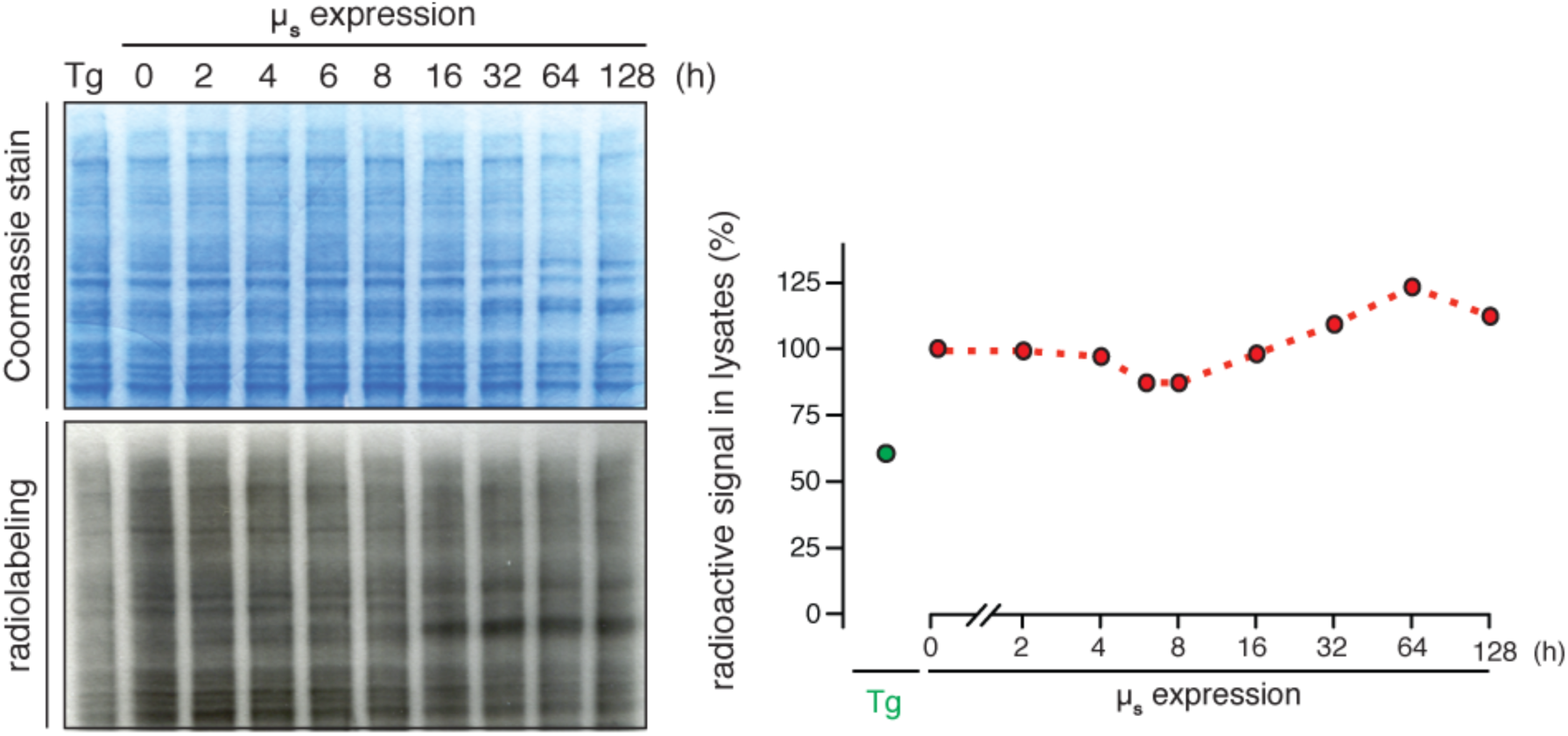
Translation is transiently attenuated upon μ_s_ overexpression to a marginal extent. HeLa-μ_s_ cells were induced with 0.5 nM Mif to express μ_s_ for the indicated times, or treated with 4 μM Tg for 45 min before they were pulse labeled for 10 min with ^35^S labeled methionine and cysteine for 10 min at the indicated times after induction, as in ***Figure 2E***. Lysates were separated by gel electrophoresis and total proteins were visualized by Coomassie, while radio-labeled proteins were detected and quantified by phosphor imaging. Labeling efficiency was compared to that in cells before induction, which was set at 100%.

